# Survivin inhibition induces PD-L1 expression through cGAS stabilization and attenuates antitumor immunity

**DOI:** 10.64898/2026.04.29.721504

**Authors:** Yun Liu, Hui Shen, Jinzhou Li, Weiqiang Yu, Tingting Yin, Xiaodong Yuan, Wenwu Luo, Yan Li, Xue Peng, Aoxing Cheng, Jing Guo, Zhenye Yang, Fazhi Yu

**Author notes:** These authors contributed equally to this work (Yun Liu, Hui Shen, Jinzhou Li, Weiqiang Yu, Tingting Yin). Correspondence to: Fazhi Yu(leading contact),; Zhenye Yang,; Jing Guo,; Hui Shen.

## Abstract

**Background:** Survivin is a mitotic regulator frequently overexpressed in human cancers and an attractive therapeutic target. However, how Survivin inhibition influences tumor immune regulation remains incompletely understood. This study aimed to investigate whether Survivin inhibition modulates antitumor immunity and to elucidate the underlying mechanisms.

**Methods:** Programmed death-ligand 1 (PD-L1) expression was evaluated in multiple tumor cell lines following pharmacological or genetic inhibition of Survivin. Mechanistic studies included RNA sequencing, immunoblotting, flow cytometry, cGAS knockdown, and NF-κB inhibition. Immune profiling was performed using CD8⁺ T-cell cytotoxicity assays, mass cytometry, flow cytometry, immunofluorescence and single-cell RNA sequencing. Clinical relevance was assessed using patient tumor specimens and public immunotherapy cohorts.

**Results:** Survivin inhibition, either by YM155 treatment or genetic depletion, increased PD-L1 expression at both mRNA and protein levels in tumor cells. Mechanistically, Survivin inhibition stabilized cyclic GMP – AMP synthase (cGAS) by reducing its ubiquitination and activated NF-κB signaling, thereby promoting transcriptional upregulation of PD-L1. Functionally, the induced PD-L1 enhanced PD-1 engagement and suppressed CD8⁺ T-cell cytotoxicity, promoting immune evasion. In immunocompetent ovarian cancer models, pharmacological inhibition of Survivin increased PD-L1 expression in tumor and immune compartments and attenuated cytotoxic immune activity. PD-L1 blockade restored antitumor immunity and significantly enhanced the therapeutic efficacy of Survivin inhibition. In addition, analyses of patient samples and public single-cell datasets revealed an inverse association between Survivin and PD-L1 expression, and high Survivin expression was associated with reduced benefit from PD-1/PD-L1 blockade.

**Conclusions:** These findings identify a Survivin–cGAS–PD-L1 axis linking mitotic stress to immune suppression and provide a mechanistic rationale for combining Survivin-targeted therapy with immune checkpoint blockade.

## Introduction

Aberrations in cell-cycle progression are a fundamental hallmark of cancer, and therapeutic targeting of mitotic regulators has long been pursued as an anticancer strategy[1–3]. Among these regulators, Survivin (encoded by *BIRC5*) is required for proper chromosome segregation and also inhibits apoptosis, thereby supporting both mitosis and tumor cell viability[4–6]. As the smallest member of the inhibitor of apoptosis protein (IAP) family, Survivin is aberrantly overexpressed in most human cancers, including ovarian cancer, but largely absent in normal adult tissues[7, 8]. This cancer-specific expression pattern has made Survivin an attractive therapeutic target, as it enables tumor suppression with minimal toxicity to normal cells[9, 10].

YM155, a small-molecule inhibitor initially developed to transcriptionally repress Survivin, induces apoptosis and inhibits tumor cell proliferation, exhibiting potent preclinical antitumor activity across various cancer types, including ovarian cancer[11–13]. Despite its cytotoxic efficacy in vitro and in immunodeficient mouse tumor models, multiple Phase I/II clinical trials of YM155 have yielded disappointing outcomes, with only limited objective responses in patients with advanced malignancies, suggesting that tumor-extrinsic resistance mechanisms may limit its clinical activity[14–17]. This discrepancy raises the possibility that Survivin inhibition may paradoxically reshape the tumor immune phenotype by modulating the expression of immune-regulatory molecules, yet whether such remodeling contributes to the limited therapeutic efficacy remains unknown.

Programmed death-ligand 1 (PD-L1) (encoded by *CD274*) -mediated immune evasion represents a major mechanism of tumor persistence under immune surveillance[18]. By binding to PD-1 on activated T cells, PD-L1 suppresses T-cell effector functions and impairs immune-mediated tumor clearance[19, 20]. Immune checkpoint blockade targeting PD-1/PD-L1 has markedly improved survival outcomes in various cancers; however, the efficacy of PD-1/PD-L1 monotherapy remains suboptimal in ovarian cancer[21, 22].

Notably, patients with PD-L1 positive status derive clinical benefit from PD-1/PD-L1 blockade[23, 24]. This observation prompts investigation into how PD-L1 expression is modulated by non-canonical stress signals such as mitotic disruption or genomic instability[25]. Canonical signaling pathways such as JAK-STAT and NF-κB are known to drive PD-L1 expression[26], yet how mitotic disruption and cell-cycle stress influence its regulation remains largely unexplored.

In this study, we identify that pharmacological inhibition of Survivin by YM155 induces PD-L1 upregulation in ovarian cancer models, which reduces T cell cytotoxicity rather than enhancing immune recognition. YM155 suppresses Survivin, stabilizes the cyclic GMP-AMP synthase (cGAS) protein, and enhances NF-κB activation, leading to transcriptional induction of PD-L1 and reduced susceptibility to CD8⁺ T-cell-mediated cytotoxicity. In vivo, YM155 increases PD-L1 levels on tumor cells and diminishes immune-mediated tumor control, whereas combination with PD-L1 blockade restores T-cell cytotoxicity and reduces tumor burden. Analyses of ovarian cancer biopsies and published patient cohorts treated with anti-PD-1/PD-L1 antibodies reveal an inverse correlation between Survivin and PD-L1 expression, and high Survivin levels are associated with shorter overall survival and lower response rates to immune checkpoint blockade. Collectively, these findings demonstrate that Survivin inhibition activates a cGAS–NF-κB–dependent PD-L1 induction program that promotes immune evasion, and provide a rationale for combining Survivin-targeted therapy with PD-1/PD-L1 blockade to improve tumor control of ovarian cancer.

## Results

### YM155 increases PD-L1 expression across multiple tumor cell types

To evaluate whether mitotic disruption influences PD-L1 expression, we treated HeyA8 ovarian cancer cells with a panel of anti-mitotic compounds and quantified PD-L1 surface levels by flow cytometry (Figure 1A). Most agents moderately increased PD-L1 expression, indicating a correlation between interference with mitotic progression and elevated PD-L1 levels (Figure 1A, B). Among these compounds, YM155 consistently produced the strongest induction, resulting in a 2- to 3-fold increase in PD-L1 surface expression relative to vehicle controls (Figure 1B). Dose-response experiments (0-20 nM) revealed a gradual, dose-dependent elevation in PD-L1 expression (Figure 1C). Additionally, time-course experiments showed a progressive increase in PD-L1 expression following YM155 treatment (Figure 1D). These findings indicate that PD-L1 expression increases in proportion to YM155 dose and duration, consistent with a regulated induction process.

**Figure 1.**
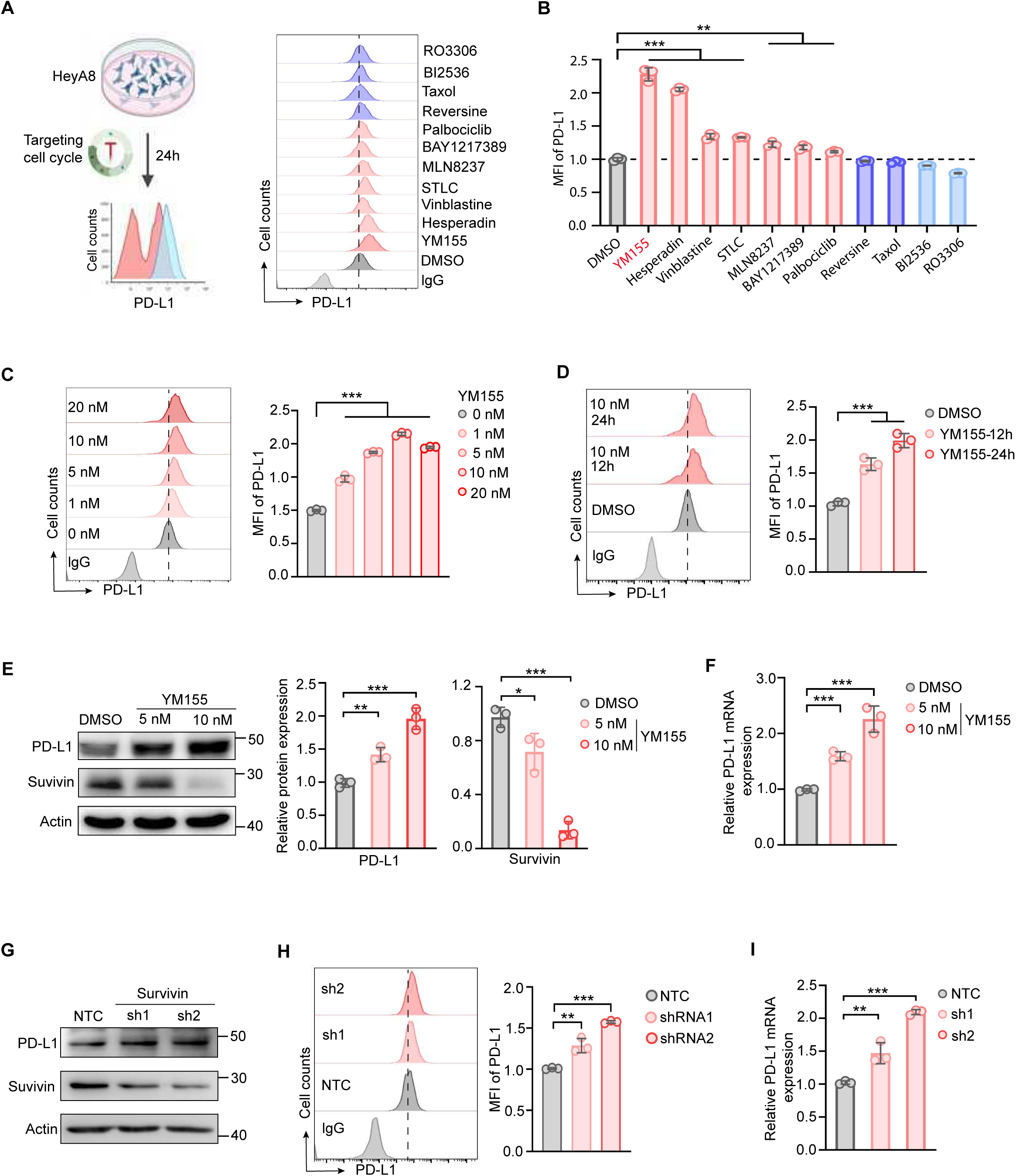
YM155 increases PD-L1 expression in HeyA8 cells. (A) Flow cytometric analysis of PD-L1 expression in HeyA8 cells treated with various anti-mitotic agents. Among all tested compounds, YM155 elicited the strongest induction of PD-L1 expression. (B) Quantification of PD-L1 surface levels from (A), presented as mean fluorescence intensity (MFI) relative to vehicle-treated controls. (C) Dose-response analysis of PD-L1 expression in HeyA8 cells treated with increasing concentrations of YM155 (1–20 nM). (D) Time-course analysis of PD-L1 expression in HeyA8 cells treated with 10 nM YM155. (E) Western blot analysis of Survivin and PD-L1 levels in HeyA8 cells treated with DMSO, 5 nM or 10 nM YM155 for 24 h. β-Actin served as the loading control. (F) qRT-PCR analysis of PD-L1 mRNA levels in HeyA8 cells treated with DMSO, 5 nM or 10 nM YM155 for 24 h. (G) Western blot analysis showing Survivin knockdown efficiency and PD-L1 expression in HeyA8 cells. (H) Flow cytometric quantification of PD-L1 surface levels in HeyA8 cells transfected with Survivin knockdown or non-targeting control (NTC) plasmids. (I) qRT-PCR analysis of PD-L1 mRNA levels in HeyA8 cells transfected with Survivin knockdown or NTC plasmids. Data are shown as mean ± SD from at least three independent experiments. p values were calculated by two-tailed Student’s t-test (ns, not significant; *p < 0.05; **p < 0.01; ***p < 0.001).

To determine whether this effect is consistent across tumor types, we extended the analysis to additional cancer cell lines. In OVCAR8 (ovarian cancer), MDA-MB-231 (triple-negative breast cancer), and BT549 (triple-negative breast cancer) cells, YM155 treatment consistently enhanced PD-L1 surface expression, suggesting that this regulatory effect is not limited to a single cell type or cancer subtype (Figure S1A-C). Western blot analysis confirmed these results, showing elevated total PD-L1 protein levels upon YM155 exposure (Figure 1E). Real-time quantitative PCR (qRT-PCR) also revealed increased *CD274* mRNA expression, suggesting that YM155 upregulates PD-L1 at the transcriptional level (Figure 1F). Together, these data show that YM155 increases PD-L1 expression in multiple tumor cell lines, supporting the generalizability of this regulatory effect.

### Survivin suppression contributes to YM155-induced PD-L1 upregulation

Because YM155 was originally developed as a Survivin suppressant, we next asked whether YM155-induced PD-L1 upregulation is linked to inhibition of Survivin[11]. Consistent with previous reports, YM155 significantly reduced Survivin protein levels in HeyA8 cells (Figure 1E). Reduction of Survivin protein was associated with increased PD-L1 expression (Figure 1E), indicating that PD-L1 upregulation may result from Survivin inhibition.

To directly test this relationship, we performed shRNA-mediated knockdown of Survivin in HeyA8 cells and assessed PD-L1 expression. Depletion of Survivin increased PD-L1 expression at both protein and mRNA levels (Figure 1G-I). These results suggest that suppression of Survivin contributes to YM155-induced PD-L1 upregulation.

To distinguish effects related to cell-cycle regulation from those related to apoptosis, we co-treated HeyA8 cells with YM155 and the pan-caspase inhibitor Z-VAD to block caspase-dependent apoptosis. Flow cytometric analysis showed that PD-L1 upregulation persisted despite the inhibition of apoptosis (Figure S1D), suggesting that YM155 primarily drives PD-L1 expression through cell-cycle regulation rather than apoptotic pathways. Persistent PD-L1 upregulation even with caspase inhibition, alongside Survivin knockdown mimicking the effect of YM155, indicates that YM155-induced PD-L1 induction is independent of caspase-mediated apoptosis.

### YM155 increases PD-L1 expression in the ovarian tumor microenvironment

To determine whether Survivin inhibition induces PD-L1 expression in vivo, we used an orthotopic ID8 ovarian cancer model in immunocompetent mice. Mice were intraperitoneally injected with ID8 cells, and three weeks later, they were treated with either YM155 or vehicle control (Figure 2A, B). Ascitic fluid was collected for mass cytometry (CyTOF) analysis, enabling high-dimensional profiling of both tumor and immune cell populations (Figure 2C). t-SNE clustering identified multiple cellular subsets, including tumor cells, macrophages, T cells, and B cells (Figure 2C, D; Figure S2A). YM155 treatment increased PD-L1 levels across several of these cellular subsets (Figure 2E; Figure S2B). Tumor cells and macrophages exhibited the most prominent PD-L1 expression increases, as reflected by elevated PD-L1 signal intensity (Figure 2E) and increased mean intensity values (Figure S2C).

**Figure 2.**
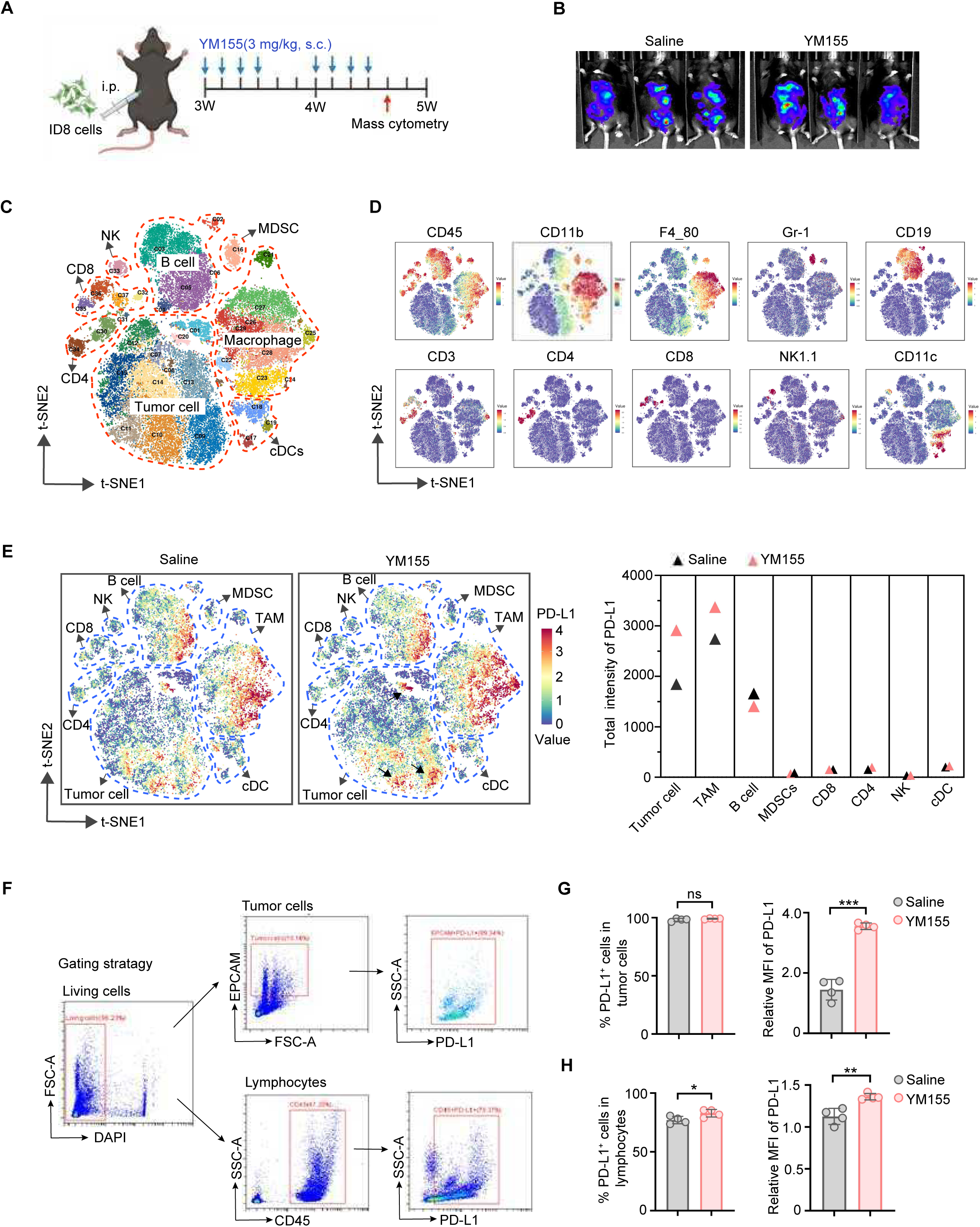
YM155 increases PD-L1 expression in tumor cells of ID8-bearing mice. (A) Schematic of the syngeneic ovarian cancer mouse model. ID8-luciferase cells were injected intraperitoneally into C57BL/6 mice, followed by treatment with saline or YM155 as indicated. Ascitic cells were collected for mass cytometry (CyTOF) analysis. (B) Tumor burden was assessed via bioluminescence imaging (n=3 mice/group). (C) t-SNE plots showing distinct cell clusters of ascitic cells from ID8-bearing mice analyzed by CyTOF. (D) Representative t-SNE plots showing lineage markers used to distinguish individual cell clusters. (E) PD-L1 expression levels across multiple cell subsets within the ascitic tumor microenvironment. (F) Flow cytometric analysis of PD-L1 expression in tumor cells and lymphocytes from saline- and YM155-treated mice. (G) Proportion and MFI quantification of PD-L1⁺ tumor cells in the ascitic microenvironment following YM155 treatment (n=4). (H) Proportion and MFI quantification of PD-L1⁺ lymphocytes in the ascitic microenvironment following YM155 treatment (n=4). Data are shown as mean ± SD from at least three independent experiments. p values were calculated by two-tailed Student’s t-test (ns, not significant; *p < 0.05; **p < 0.01; ***p < 0.001).

Furthermore, conventional flow cytometry corroborated the CyTOF data, demonstrating an increased frequency of PD-L1⁺ cells and elevated mean fluorescence intensity of PD-L1 among viable ascitic cells from YM155-treated mice (Figure S2D, E). We further examined PD-L1 expression separately in tumor cells and lymphocytes (Figure 2F). The proportion of PD-L1⁺ tumor cells remained similar between groups, but their PD-L1 mean intensity increased significantly following YM155 treatment (Figure 2G). In lymphocytes, both the percentage of PD-L1⁺ cells and their expression levels were increased after YM155 treatment (Figure 2H).

Additionally, we collected bulk tumor tissue from mice to detect PD-L1 and Survivin expression. Immunofluorescence staining showed decreased Survivin expression and increased PD-L1 expression in tumors from YM155-treated mice (Figure S2F). These data demonstrate that YM155 upregulates PD-L1 expression in vivo, especially in tumor cells, and that this response coincides with reduced Survivin expression in the tumor microenvironment.

### YM155-induced PD-L1 suppresses CD8^+^ T-cell cytotoxicity and attenuates antitumor immunity

Having established that Survivin inhibition induces PD-L1 expression in vitro and in vivo, we next asked whether the induced PD-L1 is functionally competent to suppress cytotoxic lymphocyte activity. The PD-1 binding assay showed increased PD-1 antibody binding on YM155-treated HeyA8 cells (Figure 3A, B), indicating increased surface PD-L1 with enhanced PD-1 binding capacity. Similar results were obtained in OVCAR8 and MDA-MB-231 cell lines (Figure S3A, B), suggesting that YM155 enhances functional PD-L1 expression across multiple tumor models.

**Figure 3.**
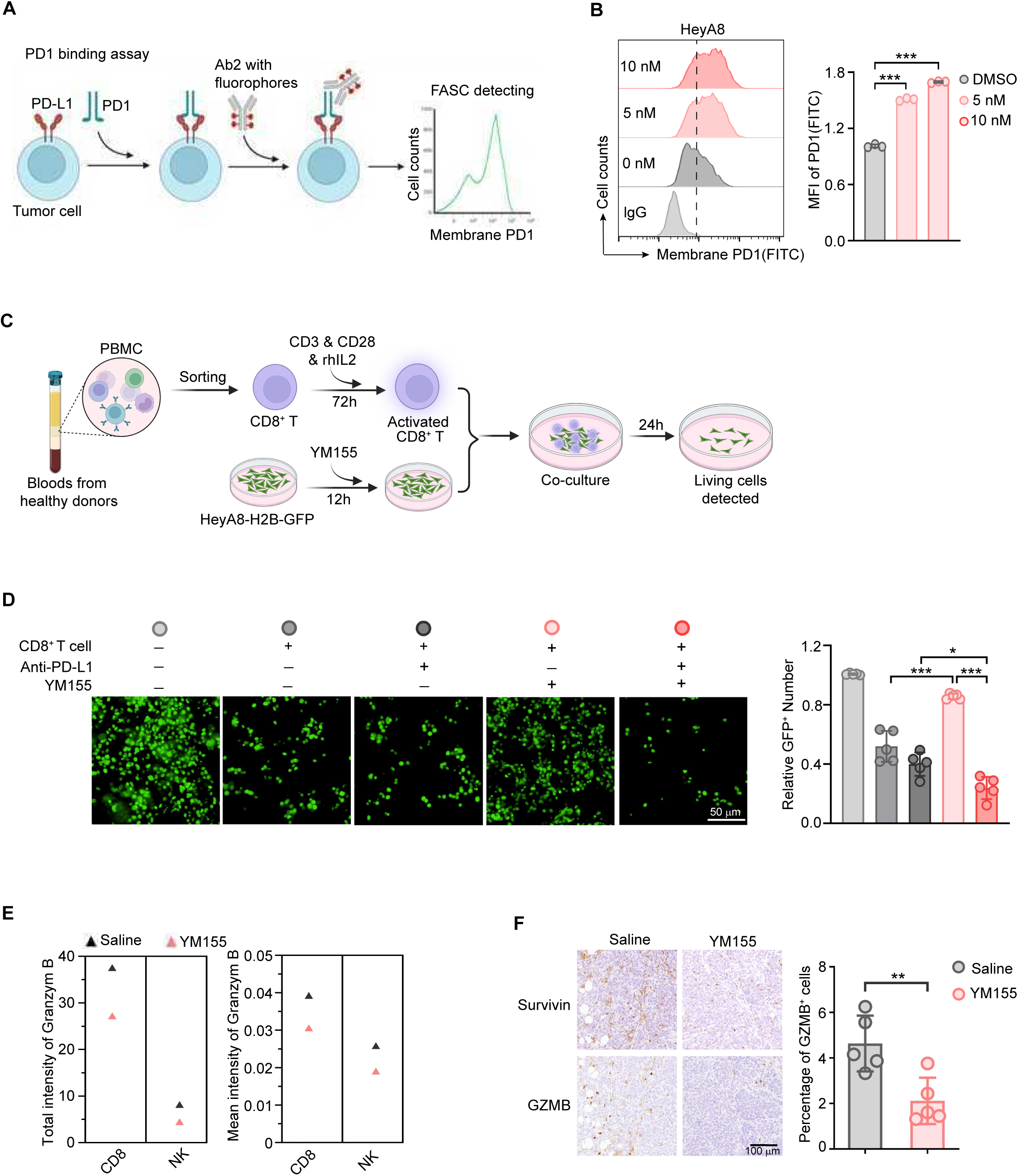
PD-L1 induced by YM155 suppresses CD8⁺T-cell cytotoxicity. (A) Schematic of the PD-1 binding assay. (B) Flow cytometric analysis of PD-1 antibody binding on the cell surface, with quantification of MFI relative to DMSO-treated controls. (C) Schematic of the co-culture cytotoxicity assay. CD8⁺ T cells were purified from healthy donor PBMCs, activated with anti-CD3/anti-CD28, and co-cultured with HeyA8-H2B-GFP tumor cells pretreated with YM155 or vehicle. (D) Representative images of the co-culture assay. The number of surviving tumor cells was quantified after co-culture. (E) CyTOF-based quantification of GZMB⁺ CD8⁺ T cells and NK cells, showing their frequency and MFI in the ascitic microenvironment following YM155 treatment. (F) Immunohistochemical staining of tumor tissues from ID8-bearing mice for Survivin and GZMB. The percentage of GZMB⁺ cells was quantified. Data are shown as mean ± SD from at least three independent experiments. p values were calculated by two-tailed Student’s t-test (*p < 0.05; **p < 0.01; ***p < 0.001).

To determine the consequence of this PD-L1 induction for antitumor immunity, we co-cultured activated human CD8^+^ T cells with HeyA8-H2B-GFP tumor cells. CD8⁺ T cells were isolated from healthy donor peripheral blood mononuclear cells (PBMCs), activated, and co-cultured with HeyA8-H2B-GFP cells (Figure 3C). YM155-treated tumor cells were more resistant to CD8⁺ T-cell-mediated killing compared to untreated cells (Figure 3D). However, this immune resistance was reversed by the addition of a PD-L1 blocking antibody, indicating that PD-L1 plays a major role in reducing T-cell cytotoxicity (Figure 3D).

In the ID8 ovarian cancer model, CyTOF analysis revealed that YM155 treatment was associated with reduced Granzyme B (GZMB) expression in CD8⁺ T cells and NK cells, indicating decreased cytolytic activity (Figure 3E). The overall cellular composition of the tumor microenvironment remained largely unchanged (Figure S3C, D), though a slight decrease in immunosuppressive cell populations (such as TAMs and MDSCs) and a mild increase in antitumor immune cells (including CD8⁺ T cells and NK cells) were observed (Figure S3E). Immunohistochemical analysis confirmed reduced Survivin and decreased numbers of GZMB⁺ cells in tumors from YM155-treated mice (Figure 3F), supporting the conclusion that YM155-induced PD-L1 dampens cytotoxic effector function. These findings demonstrate that PD-L1 induced by YM155 binds PD-1 and reduces CD8⁺ T-cell cytotoxicity, indicating that YM155-induced PD-L1 can attenuate antitumor immunity.

### PD-L1 blockade restores antitumor immunity and enhances the efficacy of YM155

Because Survivin inhibition-induced PD-L1 attenuated cytotoxic immune activity, we next investigated whether PD-L1 blockade could restore antitumor immunity and improve therapeutic efficacy. Mice bearing ID8 ovarian tumors were treated with YM155, anti-PD-L1 antibody or their combination, and tumor growth was monitored with a mouse imaging system (Figure 4A). The results showed that the combination of YM155 and anti-PD-L1 antibody resulted in significantly enhanced antitumor efficacy compared to either monotherapy or control groups (Figure 4B). Furthermore, ascitic tumor growth and ascites volume were substantially reduced (Figure 4C; Figure S4A, B), and mice receiving the combination treatment exhibited a significant improvement in overall survival (Figure 4D).

**Figure 4.**
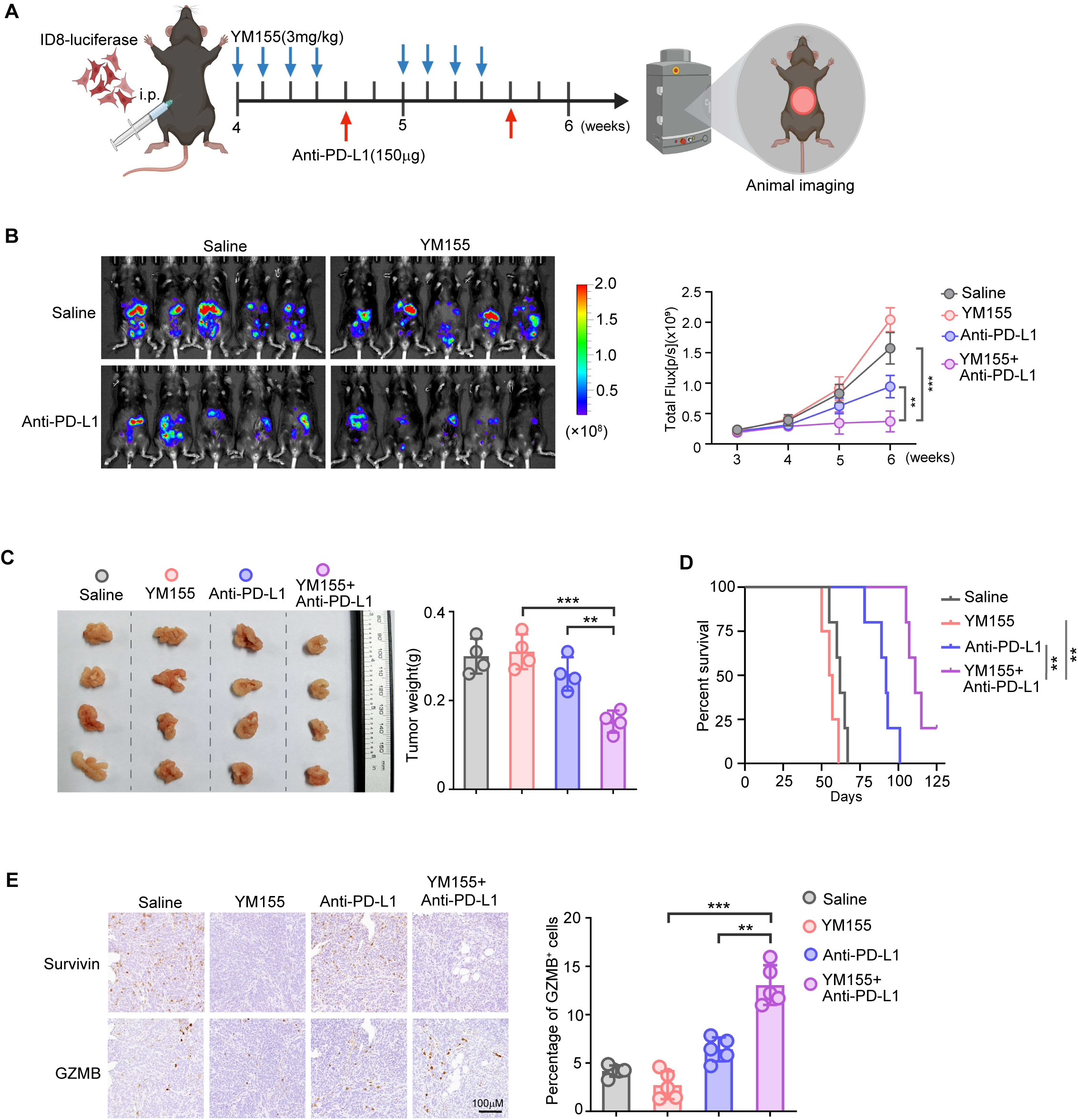
Combined YM155 and anti-PD-L1 treatment suppresses tumor growth in ID8 ovarian cancer-bearing mice. (A) Schematic of the ID8 syngeneic mouse model. ID8-luciferase cells were injected intraperitoneally into C57BL/6 mice and treated with saline, YM155, anti-PD-L1, or the combination. (B) Tumor burden was monitored via bioluminescence imaging, and tumor progression curves were plotted (n=5). (C) Representative images of ascitic tumors, with corresponding tumor weights quantified (n=4). (D) Kaplan–Meier survival curves of ID8-bearing mice treated as indicated (n=5). (E) Immunohistochemical staining of Survivin and GZMB in tumor tissues from ID8-bearing mice, with quantification of GZMB⁺ cell percentages (n=5). Data are shown as mean ± SD from at least three independent experiments. p values were calculated by two-tailed Student’s t-test (ns, not significant; **p < 0.01; ***p < 0.001).

To test whether this combination strategy extends to other tumor settings, we evaluated YM155 and anti-PD-L1 treatment in the MMTV-PyMT spontaneous breast cancer model. Consistent with findings in the ID8 model, the combination of YM155 and anti-PD-L1 treatment significantly suppressed tumor progression (Figure S4C). Immunohistochemical analysis showed reduced Survivin levels and increased infiltration of GZMB⁺ T cells in tumors treated with YM155 and anti-PD-L1 (Figure 4E).

To better recapitulate the frequent *TP53* loss-of-function in high-grade serous ovarian cancer (HGSOC) and assess whether p53 status influences treatment response, we established a syngeneic mouse tumor model by generating ID8 cells (p53 wild-type) with stable Trp53 knockout [27], and evaluated the combinatorial efficacy of YM155 and anti-PD-L1. In the p53-deficient model, the combination also reduced tumor progression compared to single-agent treatments (Figure S4D). These results show that combining Survivin inhibition with PD-L1 blockade reduces tumor burden, diminishes ascites, and prolongs survival, likely through restoring cytotoxic immune activity.

### Combined YM155 and PD-L1 blockade enhances cytotoxic immune-cell activation and infiltration

To assess the antitumor immune responses elicited by YM155 combined with anti-PD-L1 therapy, tumors from ID8-p53⁻/⁻ mouse ascites were collected and analyzed by bulk RNA sequencing. The data showed that this combination increased the expression of genes in pathways related to immune activation, including the interferon gamma response (Figure S5A). Single-cell RNA sequencing (scRNA-seq) was conducted to profile immune and tumor cell subsets within the ovarian cancer tumor microenvironment and assess treatment-induced changes associated with YM155 and/or anti-PD-L1 therapy (Figure 5A and S5B). Cell types including NK cells, monocytes, and T cells were annotated based on canonical lineage markers (Figure 5A and S5B). We found that the proportions of NK cells and T cells were higher in tumors treated with combined YM155 and anti-PD-L1 antibody than in untreated controls (Figure S5B).

**Figure 5.**
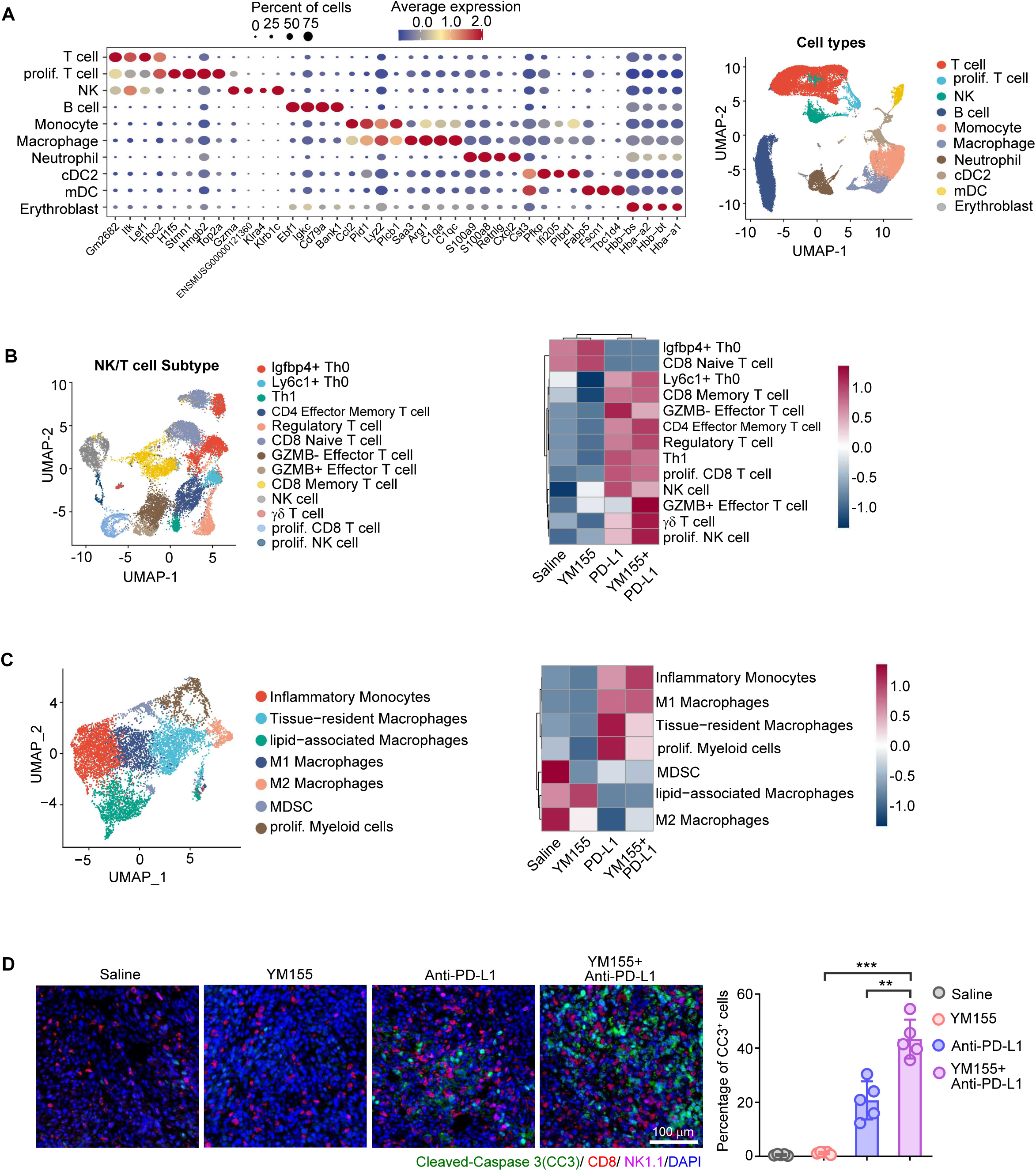
Combined YM155 and anti-PD-L1 treatment enhances activity of cytotoxic immune cells in ID8-p53⁻/⁻ mice. (A) Ascitic cells from ID8-p53⁻/⁻ mice were collected for single-cell RNA sequencing. Canonical lineage markers and t-SNE plots of identified cell clusters are shown. (B) t-SNE plots showing NK and T-cell subtypes within the tumor microenvironment. Right: Heatmap displaying the relative abundance of tumor-infiltrating lymphocyte subsets across treatment groups (Saline, YM155, PD-L1 blockade, and YM155 + PD-L1 blockade). Each row represents a distinct immune population identified by single-cell transcriptomics. (C) t-SNE plots showing myeloid cell subtypes within the ascitic tumor microenvironment, and heatmap showing their relative abundance across treatment groups. (D) Representative immunofluorescence images showing infiltration of NK cells, T cells, and cleaved-caspase-3⁺ apoptotic cells in tumor tissues from ID8-p53⁻/⁻ mice. Quantification of cleaved-caspase-3⁺ cells is shown. Data are shown as mean ± SD from at least three independent experiments. p values were calculated by two-tailed Student’s t-test (ns, not significant; **p < 0.01; ***p < 0.001).

To examine whether the cytotoxic activity of NK and T cells was also enhanced, we analyzed the composition of NK/T cell subtypes (Figure 5B and S5C). Populations associated with enhanced antitumor immune responses, such as GZMB^+^ effector T cells and proliferating NK cells, were significantly increased in the combination treatment group compared to either monotherapy or control groups (Figure 5B). In contrast, the proportion of immune-suppressive cells, including M2 macrophages and myeloid-derived suppressor cells, was reduced in the combination treatment group (Figure 5C and S5D). Immunofluorescence analysis confirmed that tumor cell death was augmented alongside increased NK and T cell infiltration within the tumor environment, indicating enhanced antitumor activity of these immune cells (Figure 5D). These results suggest that the combination of YM155 and anti-PD-L1 treatment enhances the activation and infiltration of antitumor immune cells.

### YM155 induces PD-L1 transcription through cGAS stabilization and NF-κB activation

To define the molecular mechanism by which Survivin inhibition induces PD-L1 expression, we performed bulk RNA sequencing in HeyA8 cells treated with YM155 or vehicle. YM155 treatment altered the expression of numerous genes, with 406 upregulated and 589 downregulated transcripts (Figure S6A). As expected, *BIRC5* mRNA levels were reduced, and *CD274* mRNA levels were elevated following YM155 treatment (Figure S6B). Gene set enrichment analysis (GSEA) and pathway annotation further revealed dysregulation of genes involved in cell-cycle regulation upon YM155 treatment (Figure 6A), consistent with prior findings that YM155 alters cell cycle distribution[28, 29]. Genes associated with the cytokine-cytokine receptor interaction pathway and the JAK-STAT signaling pathway were also altered by YM155 (Figure 6A). Given that PD-L1 expression can be regulated through JAK-STAT signaling pathway[30], we tested whether this pathway mediates YM155-induced PD-L1 upregulation. Treatment with the JAK1/2 inhibitor ruxolitinib failed to block PD-L1 induction by YM155 (Figure S6C, D), indicating that PD-L1 induction occurs independently of this pathway.

**Figure 6.**
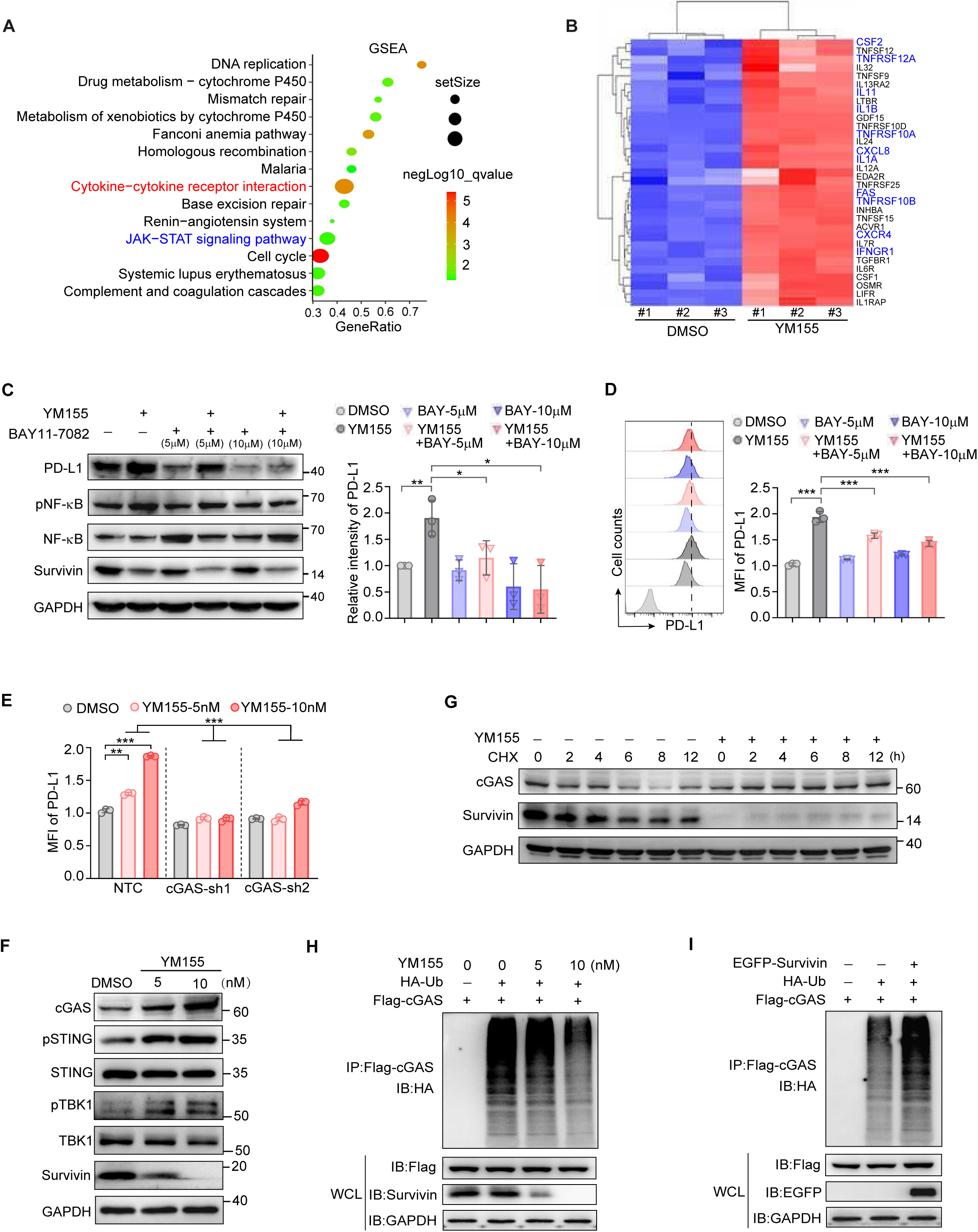
YM155 promotes PD-L1 expression through cGAS stabilization and NF-κB activation. (A) Gene Set Enrichment Analysis (GSEA) of RNA-seq data comparing YM155-treated and DMSO-treated HeyA8 cells. The top altered pathways are shown. (B) Heatmap showing differentially expressed genes in the cytokine–cytokine receptor interaction pathway. Genes involved in NF-κB signaling are highlighted in blue. (C) Left: Western blot analysis of PD-L1, pNF-κB, NF-κB, and Survivin in HeyA8 cells treated with DMSO or 10 nM YM155, in combination with 0, 5, or 10 μM BAY11-7082. GAPDH served as the loading control. Right: Quantification of protein expression levels normalized to DMSO-treated controls. (D) Left: Flow cytometric analysis of PD-L1 surface expression in HeyA8 cells treated as in (C). IgG-labeled cells served as controls. Right: Quantification of PD-L1 MFI relative to DMSO-treated controls. (E) Flow cytometric quantification of PD-L1 surface levels in HeyA8 cells transfected with cGAS knockdown or NTC plasmids, followed by treatment with DMSO, 5 nM or 10 nM YM155. Data are presented as MFI relative to DMSO-treated control cells. (F) Western blot analysis of cGAS, pSTING, STING, pTBK1, TBK1 and Survivin in HeyA8 cells treated with DMSO, 5 nM or 10 nM YM155 for 24 h. GAPDH served as the loading control. (G) Western blot analysis of cGAS and Survivin protein stability in HeyA8 cells treated with 10 nM YM155 for 12 h, followed by 100 μg/mL CHX. GAPDH served as the loading control. (H) Western blot analysis of cGAS ubiquitination in HeyA8 cells co-transfected with Flag-cGAS and HA-Ub, followed by treatment with 0, 5, or 10 nM YM155 for 24 h. cGAS was immunoprecipitated (IP) with anti-Flag antibody, and its ubiquitination level was detected by immunoblotting (IB) with anti-HA antibody. Whole cell lysates (WCL) were immunoblotting for Flag-cGAS and Survivin, with GAPDH serving as the loading control. (I) Western blot analysis of cGAS ubiquitination in HeyA8 cells co-transfected with Flag-cGAS, HA-Ub, and/or EGFP-Survivin. cGAS was immunoprecipitated (IP) with anti-Flag antibody, and its ubiquitination level was detected by immunoblotting (IB) with anti-HA antibody. WCL were immunoblotting for Flag-cGAS and EGFP-Survivin, with GAPDH serving as the loading control. Data are shown as mean ± SD from at least three independent experiments. p values were calculated by two-tailed Student’s t-test (ns, not significant; *p < 0.05; **p < 0.01; ***p < 0.001).

Notably, several inflammation-related genes within the cytokine-cytokine receptor interaction pathway were upregulated following YM155 treatment, many of which are known targets of the NF-κB signaling pathway (Figure 6B). We therefore examined whether YM155 activates NF-κB signaling as a mechanism for PD-L1 induction. Western blot and flow cytometry analysis confirmed increased levels of phosphorylated NF-κB (pNF-κB) following YM155 treatment. Importantly, co-treatment with the NF-κB inhibitor BAY11-7082 significantly suppressed YM155-induced PD-L1 expression in HeyA8 cells (Figure 6C, D), supporting a role for NF-κB in regulating this response. Similar effects were detected in MDA-MB-231 cells, indicating that NF-κB-mediated PD-L1 induction occurs in multiple cancer cell lines (Figure S6E). We further analyzed GEO datasets in which cancer cells were treated with various mitotic inhibitors. Consistent with our results in Figure 1A, targeting Aurora kinases using MLN8237, AZD1132, or VX680 induced PD-L1 expression. These drugs also upregulated genes involved in the NF-κB pathway, as revealed by GSEA analysis (Figure S6F). In contrast, BI2536 and Reversine, which failed to activate the NF-κB pathway, did not alter PD-L1 expression (Figure S6F). Together, the data indicate that PD-L1 upregulation following mitotic inhibition correlates with activation of the NF-κB pathway.

We next investigated the upstream event responsible for NF-κB activation. Given the established role of cGAS signaling in inflammatory transcriptional responses and PD-L1 regulation[31], we asked whether cGAS is required for YM155-induced PD-L1 expression. To directly test this hypothesis, we generated HeyA8 cells with stable knockdown of cGAS (Figure S6G) and examined PD-L1 expression following YM155 treatment. In cGAS-depleted cells, YM155 failed to induce PD-L1 expression, suggesting cGAS acts upstream of NF-κB-dependent PD-L1 transcription (Figure 6E). Western blot analysis showed that YM155 increased cGAS protein abundance and activated downstream cGAS–STING signaling, as evidenced by increased phosphorylation of STING and TBK1 (Figure 6F). Moreover, YM155 prolonged cGAS protein stability (Figure 6G), consistent with a potential role in mediating sustained NF-κB activation.

To investigate how YM155 enhances cGAS protein stability, we immunoprecipitated cGAS and examined its ubiquitination under DMSO or YM155 treatment. YM155 markedly reduced the ubiquitination of cGAS (Figure 6H), indicating decreased protein degradation. Because YM155 inhibits Survivin, we next tested whether Survivin modulates cGAS ubiquitination. Overexpression of Survivin increased cGAS ubiquitination (Figure 6I). Collectively, these findings support a model in which Survivin negatively regulates cGAS stability, potentially through modulation of its ubiquitination, and YM155 counteracts this effect to stabilize cGAS and drive NF-κB-dependent PD-L1 transcription.

### Survivin expression inversely correlates with PD-L1 and is associated with reduced benefit from PD-1/PD-L1 blockade

To assess the clinical significance of our findings, we examined Survivin and PD-L1 expression in paraffin-embedded ovarian tumor specimens. Immunohistochemical staining of serial sections showed that tissue areas with high Survivin staining had lower PD-L1 signal intensity, while areas with low Survivin staining exhibited higher PD-L1 levels (Figure 7A–C, Figure S7A). Quantitative image analysis showed an inverse correlation between Survivin and PD-L1 expression levels (Figure 7D, Figure S7B).

**Figure 7.**
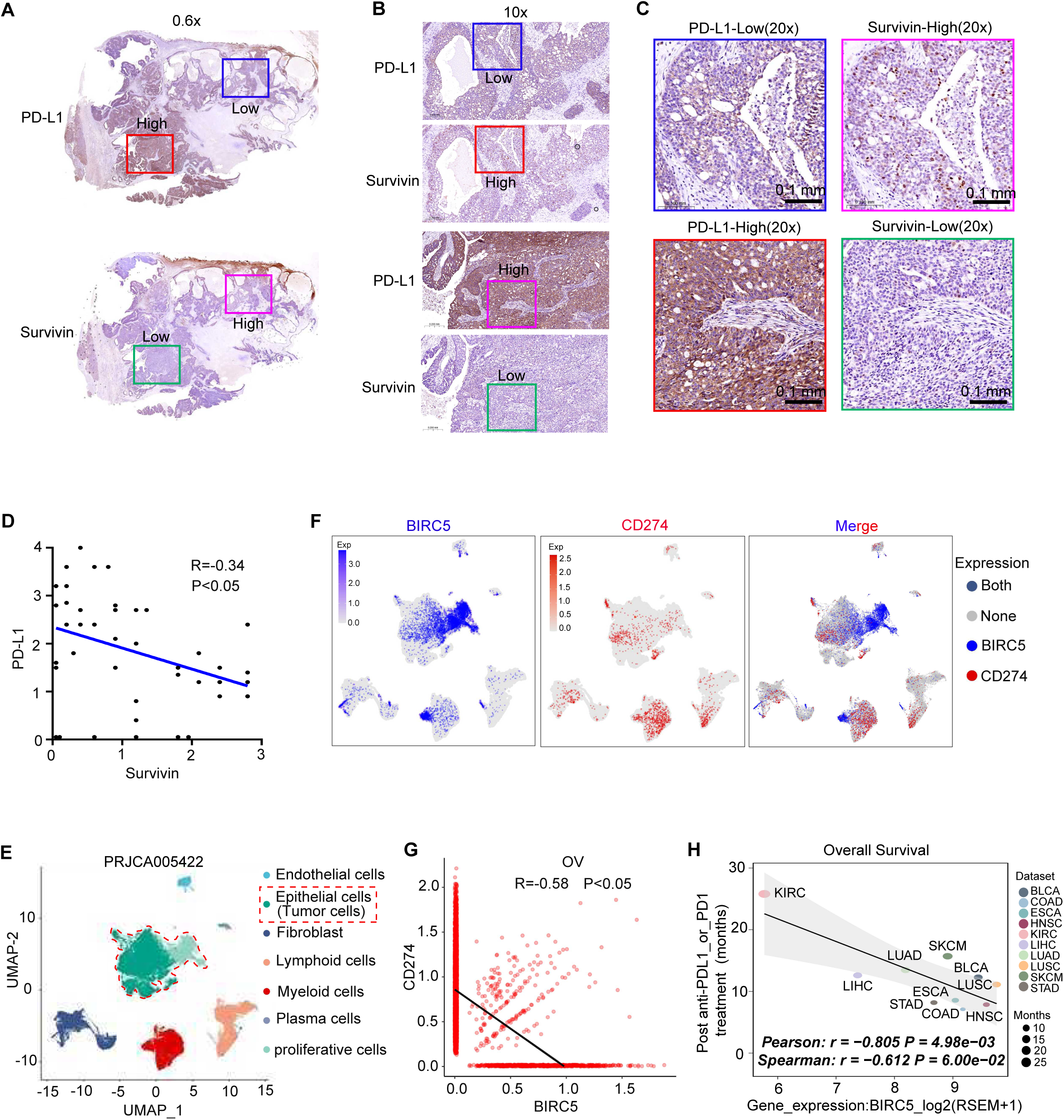
Clinical relevance of Survivin-mediated PD-L1 regulation in ovarian cancer. (A–C) Representative immunohistochemistry images of ovarian cancer patient samples stained for Survivin and PD-L1 on serial sections. Regions with high Survivin expression exhibit low PD-L1 staining (top panels), whereas low-Survivin regions show elevated PD-L1 expression (bottom panels). Scale bars = 100 μm. (B) and (C) show higher magnification views of selected areas. (D) Quantitative correlation analysis of immunostaining intensity between Survivin and PD-L1 levels. (E) Single-cell RNA sequencing (scRNA-seq) of human ovarian tumors showing UMAP clustering of identified cell types. (F) UMAP plots showing expression of BIRC5 (blue) and CD274 (red) across all tumor-infiltrating cells. The merged plot (right) indicates cells expressing BIRC5 (blue), CD274 (red), both (purple), or neither (gray). (G) Correlation analysis between BIRC5 and CD274 expression based on (F). (H) Correlation between BIRC5 expression and overall survival in patients receiving anti-PD-L1 or anti-PD-L11 therapy across multiple cancer types. Pearson or Spearman correlation coefficients indicate a statistically significant negative relationship. Statistical significance was determined by Pearson’s correlation test.

We next interrogated scRNA-seq datasets from ovarian cancer[32] and colon adenocarcinoma[33] patient samples. Cell populations were classified based on lineage markers (Figure S7C, D and 7E). In epithelial cells from both ovarian and colon cancer samples, subpopulations with high *BIRC5* expression exhibited lower *CD274* expression, based on single-cell transcriptomic data (Figure 7F; Figure S7E). Overall, across single-cell populations, cells expressing high levels of *BIRC5* exhibited reduced *CD274* expression, whereas cells with low *BIRC5* levels had elevated *CD274* (Figure 7G; Figure S7E). These patient-derived data are consistent with our in vitro and in vivo observations of inverse Survivin–PD-L1 expression.

Finally, we analyzed published cohorts of patients with advanced cancers treated with anti-PD-1 or anti-PD-L1 antibodies. The results showed that higher *BIRC5* expression was associated with shorter overall survival and reduced response rates in patients treated with anti-PD-L1 or anti-PD-1 agents (Figure 7H; Figure S7F). Taken together, these results suggest that Survivin and PD-L1 may represent potential correlates of immune phenotypes and therapeutic outcomes.

## Discussion

In this study, we identify a mechanism linking mitotic regulation to tumor immune evasion, which may partly explain why Survivin inhibition has not yet produced sustained clinical responses. We demonstrate that pharmacological inhibition of Survivin by YM155 induces PD-L1 expression across multiple tumor types, thereby promoting immune escape rather than immune activation. This PD-L1 upregulation is mediated by activation of the cGAS–NF-κB pathway. YM155 stabilizes cGAS protein, enhances NF-κB phosphorylation, and drives PD-L1 transcription, producing functionally active PD-L1 that suppresses cytotoxic T-cell activity. In mouse models, YM155 increases PD-L1 levels in tumor cells and macrophages, thereby impairing immune cell-mediated tumor clearance. Importantly, YM155-induced immune escape can be overcome by combining with anti-PD-L1 antibody, which restores cytotoxic activity of CD8⁺ T cells and NK cells and achieves potent tumor suppression compared to either monotherapy alone. Clinically, Survivin expression inversely correlates with PD-L1 levels and is associated with poor response to anti-PD-1/PD-L1 immunotherapy, highlighting the translational relevance of this immune-regulatory pathway. These findings establish a Survivin–cGAS–PD-L1 axis that links mitotic stress to adaptive immune suppression and provide a mechanistic rationale for combining Survivin-targeted therapy with immune checkpoint blockade (Figure 8).

**Figure 8.**
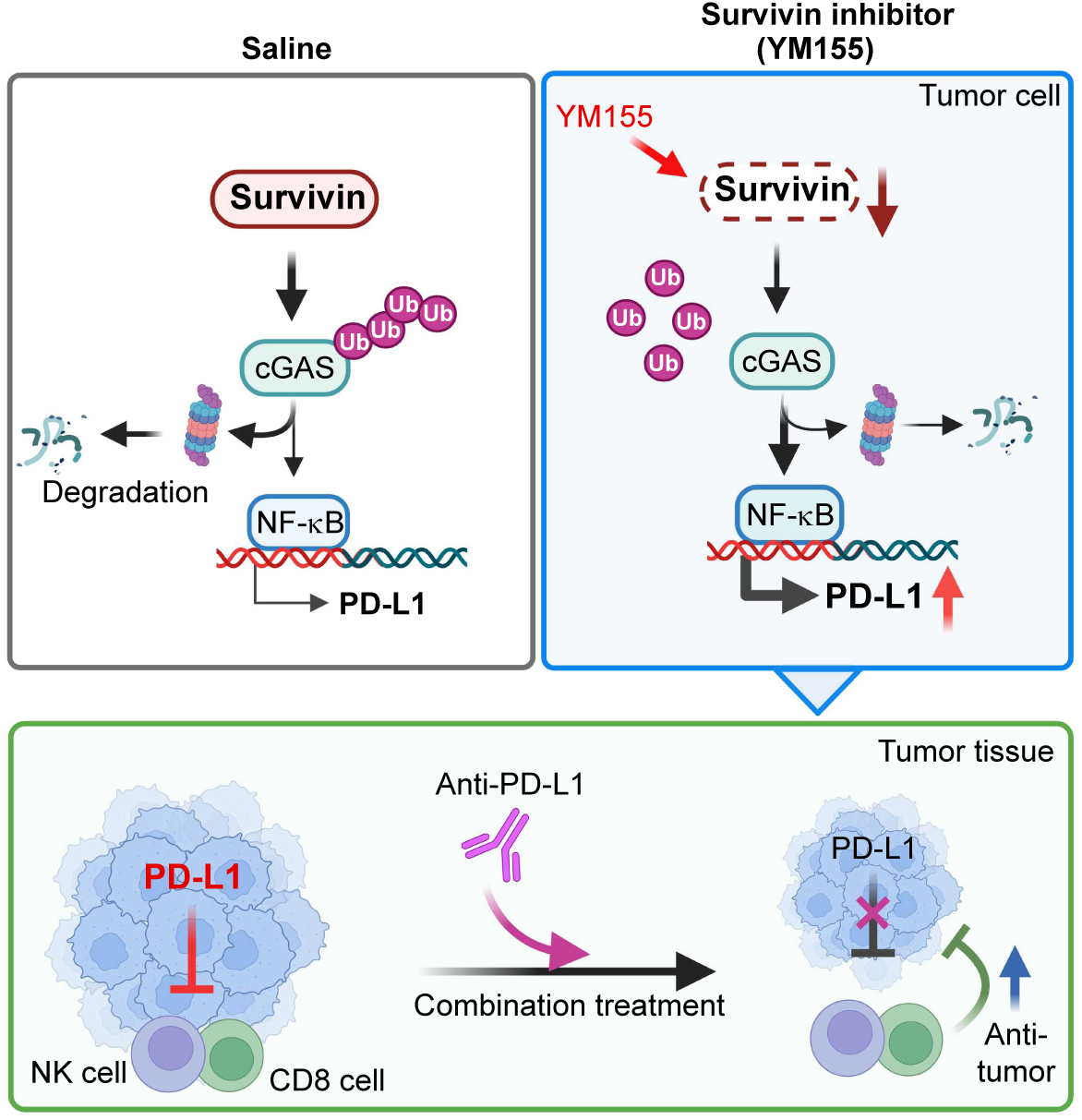
Graphic summary, YM155 targets Survivin to modulate PD-L1 expression and sensitize tumors to immune checkpoint blockade. Inhibition of the key cell cycle regulator Survivin by YM155 upregulates PD-L1 expression through activation of the cGAS–NF-κB signaling pathway, thereby transiently dampening cytotoxic immune responses. This immunosuppressive effect can be effectively neutralized by PD-L1 blockade. Consequently, combined YM155 and anti-PD-L1 therapy suppresses tumor growth by simultaneously restraining tumor proliferation and reactivating NK and CD8⁺ T-cell cytotoxicity. Collectively, these findings highlight the potential of combinatorial strategies integrating Survivin inhibition with immune checkpoint blockade to enhance antitumor immunity and overcome resistance to immunotherapy.

A central observation of our study is that YM155-mediated cGAS stabilization constitutes a key event in this axis. Under basal conditions, cGAS levels are tightly controlled by the ubiquitin-proteasome system[34, 35]. Recent work has demonstrated that the deubiquitinase TRABID stabilizes Survivin and Aurora B during mitosis. Inhibition of TRABID protects cGAS from autophagic degradation, thereby activating the cGAS – STING pathway and sensitizing tumors to anti-PD-1 therapy[36]. Extending this paradigm, our findings indicate that Survivin inhibition by YM155 triggers mitotic stress that stabilizes cGAS via ubiquitination and activates NF-κB-dependent PD-L1 transcription. Thus, altered ubiquitin-mediated turnover of cGAS emerges as a key determinant of the immunomodulatory effects of Survivin-targeted therapy.

Some cell-cycle regulators are increasingly recognized as critical modulators of tumor immunity beyond their classical role in controlling cell proliferation[37, 38]. Studies have shown that cell-cycle proteins modulate the tumor immune microenvironment by regulating the functional status of immune cells, thereby shaping antitumor immune responses[39]. This underscores the potential of cell-cycle regulators as therapeutic targets and facilitates better responses to immunotherapy. For example, CDK4/6 inhibitors, recognized for their role in arresting tumor cell-cycle progression, exert profound modulatory effects on the tumor immune landscape which reprograms the tumor immune microenvironment and enhances responsiveness to checkpoint blockade[40–42]. Recently, inhibition of Aurora A kinase, a mitotic serine/threonine-protein kinase, was reported to upregulate PD-L1 expression via cGAS dephosphorylation and STING/NF-κB pathway, ultimately compromising their own antitumor efficacy[43]. However, unlike these agents that ultimately potentiate antitumor immunity, Survivin inhibition triggers PD-L1 induction in tumor cells and suppresses cytotoxic T-cell activity that facilitates immune escape. This distinct outcome reveals a stress-adaptive mechanism through which tumor cells tolerate mitotic disruption by engaging immune checkpoint pathways. Notably, PD-L1 upregulation persists even after apoptosis blockade, suggesting that mitotic perturbation, rather than cell-death signaling, predominantly drives this process. Our data show that mitotic stress from Survivin inhibition activates the cGAS–NF-κB pathway, leading to PD-L1 transcription and functional immune suppression, thereby directly linking cell-cycle disruption with immune evasion mechanisms.

Survivin, a member of the IAP family and a regulator of mitotic progression, is aberrantly overexpressed in most human malignancies and associated with therapy resistance and poor prognosis [4, 12, 17]. Previous studies have correlated Survivin expression with immunosuppressive cell infiltration and immune checkpoint gene signatures across cancers[44, 45], suggesting its potential as an immunotherapeutic target. However, the underlying mechanism has not been fully elucidated. Despite the pronounced antitumor efficacy of the Survivin inhibitor YM155 in preclinical models, its clinical performance has been unsatisfactory with limited single-agent activity, partly attributed to incomplete understanding of its mechanism of action[17]. Recent studies have proposed YM155 can exert cytotoxic activities through multiple mechanisms including inhibiting DNA topoisomerase, ATR kinase and deubiquitylation[46, 47]. Here, we uncover an unanticipated biological consequence of Survivin inhibition: YM155 not only disrupts mitosis but also elicits adaptive immune suppression through PD-L1 upregulation via the cGAS–NF-κB pathway. Specifically, YM155 induces PD-L1 expression concomitantly with a reduction in T-cell cytotoxicity, compromising its antitumor efficacy by creating an immunosuppressive microenvironment. This effect provides a potential explanation for the suboptimal clinical efficacy observed with YM155 monotherapy. On the other hand, tumors with intrinsically low PD-L1 expression typically display limited sensitivity to PD-L1/PD-1 blockade, while YM155-induced upregulation of PD-L1 can enhance the therapeutic efficacy of this immune checkpoint blockade, establishing a rationale for combining Survivin inhibition with PD-1/PD-L1 blockade therapy. In our preclinical models, YM155 and anti-PD-L1 treatment cooperate to enhance the cytotoxic activity and infiltration of CD8⁺ T-cells and NK cells, reduce tumor burden, and thus overcome YM155-induced immune suppression, partially remodeling the tumor microenvironment toward a more immunologically active phenotype. These findings indicate that Survivin suppression activates a cGAS– NF-κB dependent PD-L1 induction program and reveal a role for Survivin in linking mitotic control with immune modulation.

Although our data delineate a functional Survivin–cGAS–NF-κB-PD-L1 signaling pathway, several questions remain unresolved. The molecular events through which Survivin depletion increases cGAS stability remain to be defined and may involve mitosis-associated DNA leakage, micronuclei formation, or altered proteasomal turnover. Survivin is a proliferation marker, and its association with immunotherapy outcomes may therefore not be entirely specific to the mechanism defined here. In addition, although YM155 and PD-L1 blockade cooperate to reduce tumor burden in our models, clinical translation will require optimization of dose, timing, and treatment sequence to balance cytotoxic and immune-modulatory effects. It will also be important to determine whether inhibitors of other mitotic regulators (e.g., PLK1 or CDC20) activate cGAS– NF-κB signaling and increase PD-L1 expression, thereby establishing whether mitotic control broadly influences tumor immunogenicity. Overall, our findings indicate that mitotic disruption can induce a stress-adaptation program that elevates PD-L1 through cGAS–NF-κB signaling and support further investigation of combined Survivin suppression and PD-1/PD-L1 blockade in tumors with limited responsiveness to immunotherapy.

## Materials and Methods

### Cell Lines and Reagents

HeyA8 and OVCAR8 ovarian cancer cells, as well as MDA-MB-231 and BT549 triple-negative breast cancer cells, were obtained from the American Type Culture Collection (ATCC) and maintained in Dulbecco’s Modified Eagle Medium (DMEM; Gibco) supplemented with 10% fetal bovine serum (FBS; Gibco) and 1% penicillin–streptomycin (Gibco). All cell lines were cultured at 37 °C in a humidified atmosphere containing 5% CO₂ and were routinely tested for Mycoplasma contamination using a PCR-based assay prior to experimentation. YM155, BAY11-7082, Z-VAD-FMK, and other anti-mitotic agents were purchased from Selleck Chemicals (Houston, TX, USA) or MedChemExpress (Monmouth Junction, NJ, USA). All compounds were dissolved in dimethyl sulfoxide (DMSO) or sterile saline according to the manufacturer’s instructions and stored as recommended.

### Lentiviral shRNA Transduction and Stable Cell Line Generation

Short hairpin RNAs (shRNAs) targeting specific genes or a non-targeting control sequence were cloned into lentiviral expression vectors. Lentiviral particles were produced by co-transfecting HEK293T cells with the shRNA construct and packaging plasmids (psPAX2 and pMD2.G) using polyethyleneimine (PEI; Polysciences, USA). Viral supernatants were collected 48 hours after transfection, filtered through 0.45 μm non-pyrogenic filters (Merck Millipore, USA), and supplemented with 8 μg/mL polybrene (Sigma-Aldrich, USA). Target cells (1 × 10⁵ per 35 mm dish) were infected with the viral supernatant for 24 hours, followed by replacement with fresh complete medium. Stably transduced cells were selected with puromycin (1-2 μg/mL) for at least 5 days. Knockdown efficiency was confirmed by Western blot analysis before use in subsequent experiments.

### Flow Cytometry

For surface PD-L1 detection, cells were gently detached with trypsin, washed with phosphate-buffered saline (PBS), and resuspended in flow buffer (PBS containing 2% fetal bovine serum). Cells were stained with fluorophore-conjugated anti-PD-L1 antibodies or the corresponding isotype controls for 30 minutes on ice. After washing, samples were analyzed on a CytoFLEX flow cytometer (Beckman Coulter), and data were processed using FlowJo software (TreeStar, Ashland, OR, USA). For PD-1 binding assays, cells were incubated with PD-1-Fc fusion proteins (R&D Systems) or PD-1 antibodies at 4 °C, followed by staining with fluorophore-conjugated secondary antibodies specific for the Fc domain. For profiling tumor-infiltrating immune cells, ascitic fluid was collected and treated with Red Blood Cell Lysis Buffer (Beyotime Biotechnology) at 4 °C for 10-15 minutes to remove erythrocytes. After washing, cells were counted using an automated cell counter (Countstar, Shanghai, China). A total of 1 × 10⁶ cells were blocked with FcR Blocking Reagent (BioLegend) or 10% mouse serum, followed by incubation with surface marker antibodies (BioLegend). Flow cytometric acquisition was performed on the CytoFLEX flow cytometer (Beckman Coulter).

### Western Blot

Cells were seeded into culture dishes 12 h prior to treatment. After treatment, culture media were aspirated, and cells were washed three times with PBS. For protein extraction, cells were lysed on ice for 30 min using lysis buffer (50 mM Tris-HCl, pH 7.4, 250 mM NaCl, 1 mM EDTA, 50 mM NaF, and 0.5% Triton X-100) supplemented with protease and phosphatase inhibitors. Lysates were centrifuged at 12,000 rpm for 15 min at 4°C, and supernatants were collected. Protein concentrations were determined using the Bradford assay (Sangon Biotech, Shanghai, China). Equal amounts of total protein (10-20 μg) were separated by SDS-PAGE and transferred onto PVDF membranes (Millipore). Membranes were blocked with 5% non-fat milk and incubated overnight at 4°C with primary antibodies. After washing, membranes were incubated with HRP-conjugated secondary antibodies and developed using enhanced chemiluminescent substrates (Western Lightning Chemiluminescence Reagent Plus, Advansta). Densitometric analysis was performed using ImageJ software.

### RNA-Seq and pathway enrichment analysis

For HeyA8 cells, total RNA (1 μg per sample) was used for mRNA library preparation, which was conducted by Genewiz (Suzhou, China). Poly(A)^+^ RNA was enriched using Oligo(dT) beads, fragmented with divalent cations at high temperature, and subjected to first- and second-strand cDNA synthesis using random hexamer primers. The resulting double-stranded cDNA was end-repaired, A-tailed, and ligated with sequencing adaptors, followed by size selection using DNA Clean Beads and PCR amplification with indexed primers. Libraries were sequenced on an Illumina HiSeq, NovaSeq, or MGI2000 platform (2 × 150 bp). Raw reads were processed with Cutadapt (v1.9.1) to remove adapters and low-quality reads (Phred < 20), then aligned to the reference genome using HISAT2 (v2.2.1). Gene expression was quantified using HTSeq (v0.6.1), and differentially expressed genes (DEGs) were identified using DESeq2 (Padj ≤ 0.05). GO and KEGG analyses were performed using GOseq (v1.34.1) and in-house scripts.

For ID8-derived tumor tissues, total RNA was extracted using TRIzol reagent (Invitrogen, USA). RNA purity and integrity were assessed with a NanoDrop 2000 and Agilent 2100 Bioanalyzer, respectively. Library construction was performed using the VAHTS Universal V10 RNA-seq Library Prep Kit (Vazyme), and sequencing and downstream analysis were completed by OE Biotech (Shanghai, China) using the Illumina NovaSeq 6000 platform. Clean reads were processed with fastp, mapped with HISAT2, and quantified with HTSeq-count. DEGs were identified using DESeq2 (Q < 0.05, fold change >2 or <0.5). PCA, hierarchical clustering, and heatmaps were performed in R (v3.2.0). GO, KEGG, Reactome, and WikiPathways enrichment analyses were conducted using hypergeometric testing and visualized with bar plots, chord diagrams, and bubble charts. GSEA was conducted using the GSEA software with predefined gene sets.

### Gene expression analysis by qRT-PCR

Total RNA was extracted using TRIzol reagent (Invitrogen) according to the manufacturer’s instructions. Residual genomic DNA was removed by DNase I treatment. Complementary DNA (cDNA) was synthesized from 1 μg of total RNA using a reverse transcription kit (Vazyme, China). Quantitative real-time PCR (qRT-PCR) was performed using SYBR® Premix (Vazyme, China) on a PikoReal 96 real-time PCR system (Thermo Scientific, MA, USA). All reactions were carried out in triplicate. Relative gene expression levels were calculated using the 2^−ΔΔCT^ method, with β-actin serving as the internal control. Primers were designed using PrimerBank or obtained from OriGene (OriGene Technologies, Inc.), and their specificity was confirmed via NCBI Primer-BLAST. All primers were synthesized by SANGON Biotech (Shanghai, China).

### T Cell Cytotoxicity Assays

Peripheral blood mononuclear cells (PBMCs) from healthy donors were obtained via Ficoll-Paque density gradient separation. CD8^+^ T cells were purified by negative selection (Miltenyi Biotec) and stimulated with anti-CD3/anti-CD28 beads (Thermo Fisher) in RPMI 1640 with 10% FBS for 72 hours. Activated T cells were co-cultured with HeyA8-H2B-GFP cells at a 5:1 effector-to-target ratio. Tumor cell killing was assessed by measuring GFP^+^ tumor cell viability via flow cytometry or fluorescence microscopy over 24 hours. In some experiments, anti-PD-L1 blocking antibodies were added to the co-culture to assess rescue from immune-mediated killing.

### In vivo mouse tumor models

All animal experiments were conducted in accordance with institutional guidelines and were approved by the Institutional Animal Care and Use Committee (IACUC) of the First Affiliated Hospital of the University of Science and Technology of China. Female C57BL/6 mice (6-8 weeks old; SLAC Laboratory Animal) were intraperitoneally (i.p.) injected with either 5 × 10⁶ ID8-p53-luciferase wild-type cells or 5 × 10⁵ ID8-p53⁻/⁻-luciferase cells. Following tumor establishment (3 weeks for wild-type and 1 week for p53-deficient cells), mice were randomly assigned to receive one of the following treatments: vehicle control, YM155 (3 mg/kg, subcutaneous injection), anti-PD-L1 antibody (clone 10F.9G2, Bio X Cell; 150 μg dissolved in PBS, administered i.p.), or a combination of both agents. Each experimental group contained 10 mice, with 5 mice pre-assigned to survival analysis and another 5 mice for tumor phenotype analysis. Tumor burden was evaluated using bioluminescence imaging. At the experimental endpoint, ascites volume was measured, and peritoneal washings were collected for flow cytometry, CyTOF and/or scRNA-seq analysis. No animals or data points were excluded from the analysis.

### Mass Cytometry (CyTOF) and t-SNE Analysis

Mass cytometry was performed using a Helios™ mass cytometer (Fluidigm, USA), as previously described [48], with modifications for ascites samples. Fresh ascites were collected from ID8 tumor-bearing mice, incubated with Red Blood Cell Lysis Buffer (Beyotime Biotechnology, China) for 10-15 minutes at 4°C to lyse erythrocytes, and then filtered through a 40 μm cell strainer. Cell suspensions were washed, counted, and resuspended at the appropriate concentration for downstream staining. A panel of about 40 metal-conjugated antibodies targeting surface and intracellular markers was used for cell phenotyping and functional analysis. Antibodies were either purchased pre-conjugated or labeled in-house using the Maxpar® Antibody Labeling Kit (Fluidigm), following the manufacturer’s instructions. Cell suspensions were incubated with surface marker antibodies, followed by fixation, permeabilization, and intracellular staining using a commercial Maxpar staining kit (Fluidigm). Stained cells were analyzed on the Helios™ mass cytometry platform (Fluidigm). During acquisition, EQ™ Four Element Calibration Beads (Fluidigm) were added for signal normalization. All data acquisition and primary quality control were conducted by PLTTech Inc. (Hangzhou, China). Raw CyTOF data were normalized using the Helios normalization algorithm, and downstream analyses were performed using Cytobank (https://www.cytobank.org/) and R-based packages such as cytofkit. Dimensionality reduction was performed using t-distributed stochastic neighbor embedding (t-SNE), and unsupervised clustering was conducted using the PhenoGraph algorithm to identify phenotypically distinct cell subsets.

### Immunofluorescence

Immunofluorescence staining was performed as previously described[49], with modifications for tumor tissue processing. Mouse tumor tissues were fixed in 4% formaldehyde for 14 hours at 4°C, dehydrated in 30% sucrose for 24 hours at 4°C, embedded in OCT compound, and stored at −80°C until sectioning. Frozen sections were permeabilized with 0.2% Triton X-100 in PHEM buffer and blocked with 1% bovine serum albumin (BSA) in TBST for 30 minutes at room temperature. Sections were then incubated with primary antibodies for 2 hours at room temperature, washed three times in TBST, and incubated with species-appropriate fluorophore-conjugated secondary antibodies for 1 hour. Nuclei were counterstained with 4’,6’-diamidino-2-phenylindole (DAPI) for 3 minutes. Coverslips were mounted using ProLong™ Gold Antifade Mountant (Thermo Fisher Scientific or Sigma-Aldrich). Whole-slide fluorescence images were acquired at 20× magnification using a Pannoramic 250 Flash III scanner (3DHISTECH). Multichannel images were visualized and reviewed using CaseViewer software. Quantitative analysis of fluorescence intensity and co-localization, where applicable, was performed using ImageJ.

### Immunohistochemistry

Formalin-fixed, paraffin-embedded tumor sections were cut at 5 µm thickness. Slides were deparaffinized, rehydrated, and subjected to heat-mediated antigen retrieval in citrate buffer (pH 6.0). Endogenous peroxidase activity was quenched by incubation in 3% hydrogen peroxide. Sections were blocked in 5% normal serum and incubated overnight with primary antibodies at 4°C. After washing, HRP-conjugated secondary antibodies were applied, and DAB was used as the chromogenic substrate. Slides were counterstained with hematoxylin and mounted. Stained slides were scanned using a whole slide scanner (Pannoramic MIDI, 3DHISTECH) at 20 × magnification to generate high-resolution digital images. Whole-slide images were viewed and analyzed using CaseViewer (3DHISTECH) software. Quantification of staining intensity and positive cell proportions was performed either manually or using image analysis algorithms as specified.

### Single-Cell RNA Sequencing and data preprocessing

Fresh ascites samples were collected from ID8 tumor-bearing mice and incubated with Red Blood Cell Lysis Buffer (Beyotime Biotechnology) for 10-15 minutes at 4°C to lyse red blood cells. Single-cell suspensions were filtered through a 40 μm strainer, counted, and adjusted to a concentration of approximately 700-1,200 cells/μL. The cell suspensions were immediately loaded into a 10x Genomics Chromium Controller for single-cell encapsulation and cDNA library preparation using the Chromium Next GEM Single Cell 3’Reagent Kits v3.1 (10x Genomics), according to the manufacturer’s protocol. Single-cell RNA sequencing was performed by OE Biotech Co., Ltd.

Libraries were sequenced on an Illumina NovaSeq 6000 platform to generate paired-end 150 bp reads. Raw sequencing data were processed using the Cell Ranger software pipeline (version 5.0.0) provided by 10x Genomics. This pipeline was used to demultiplex cellular barcodes, align reads to the mouse reference genome (mm10) using STAR, and generate a gene-by-cell expression matrix based on unique molecular identifier (UMI) counts.

Downstream analysis was conducted using the Seurat R package (version 3.1.1). Raw UMI count matrices were normalized using the NormalizeData function with the LogNormalize method, which scales gene expression by total expression per cell and log-transforms the result. Cells were filtered to exclude low-quality or potential doublet cells based on the following criteria: (i) fewer than 200 detected genes, (ii) fewer than 1,000 UMIs, (iii) log10GenesPerUMI < 0.7, (iv) >10% of transcripts mapped to mitochondrial genes, and (v) >5% mapped to hemoglobin genes. Dimensionality reduction was performed using Principal Component Analysis (PCA) followed by Uniform Manifold Approximation and Projection (UMAP). Cell clustering was carried out using the FindClusters function, and clusters were annotated based on canonical marker genes.

To explore the relationship between Survivin (BIRC5) and PD-L1 (CD274) expression, cells were stratified into BIRC5^high^ and BIRC5^low^ subpopulations based on normalized expression thresholds. PD-L1 expression was then visualized across these subpopulations using UMAP to assess the potential association between Survivin and PD-L1 expression levels.

### Statistical Analysis

All in vitro experiments were performed in at least three independent biological replicates unless otherwise specified. Data visualization and statistical analyses were conducted using GraphPad Prism or R. For group comparisons, two-tailed Student’s t-tests were used, or one-way ANOVA followed by Tukey’s post hoc test for multiple comparisons. Survival analyses (Kaplan-Meier curves) were evaluated using the log-rank (Mantel-Cox) test. A p-value of < 0.05 was considered statistically significant. Exact p-values and confidence intervals are provided in the figure legends where applicable.

## Acknowledgments

This work was supported by the National Natural Science Foundation of China (32000492), the Natural Science Foundation of Anhui Province (2508085MH187) to Fazhi Yu, and the Fundamental Research Funds for the Central Universities (YD9110002076) to Yun Liu. We thank Dr. Qinglei Gao at Huazhong University of Science for providing cell lines.

## Data and code availability

RNA sequencing data, including tumor cell line, single-cell, and bulk tumor datasets generated in this study, have been deposited in the NCBI Gene Expression Omnibus (GEO) under accession number PRJNA1344606.

## Code Availability

All codes used for data analysis are available from the corresponding author upon reasonable request.

## Ethics approval statement and Patient Consent statement

All animal experiments were performed in accordance with institutional guidelines and approved by the Animal Ethics Committee of the First Affiliated Hospital of the University of Science and Technology of China (Approval No. 2025-N(A)-016). The study involving human samples was approved by the Ethics Committee of the First Affiliated Hospital of the University of Science and Technology of China (Approval No. 2025KY-388), and written informed consent was obtained from all participants.

## Author Contributions

F.Z.Y. designed the study. F.Z.Y., H.S., and J.Z.L. performed the mechanistic experiments. W.Q.Y. analyzed the sequencing data. F.Z.Y., Y.L. (Yun Liu), and X.D.Y. conducted the in vivo assays. F.Z.Y., Y.L. (Yun Liu), H.S., and J.Z.L. performed the in vitro assays. W.W.L. carried out the immunohistochemical analyses and prepared clinical tissue samples. F.Z.Y., T.T.Y., Y.L. (Yan Li) and X.P. analyzed the clinical data. F.Z.Y. and Y.L. (Yun Liu) wrote the original draft. F.Z.Y., Y.L. (Yun Liu), H.S., T.T.Y., A.X.C., J.G., and Z.Y.Y. reviewed and edited the manuscript. F.Z.Y., Z.Y.Y., J.G., and H.S. supervised the study.

