## Supplementary figures and images for "Survivin inhibition induces PD-L1 expression through cGAS stabilization and attenuates antitumor immunity"

### Supplemental figure

**Figure S1**

A

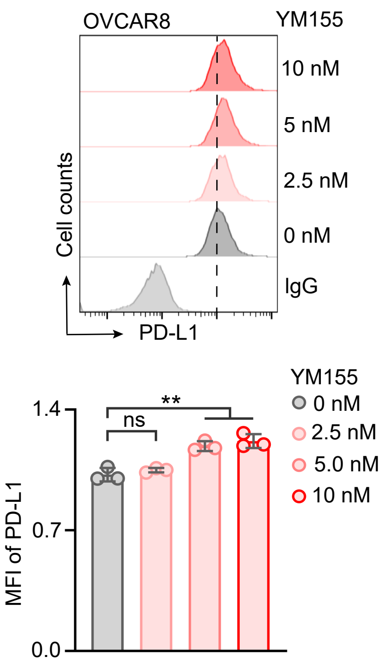


B

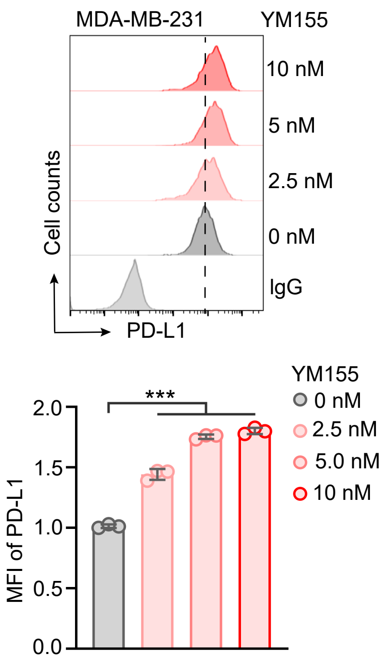


C

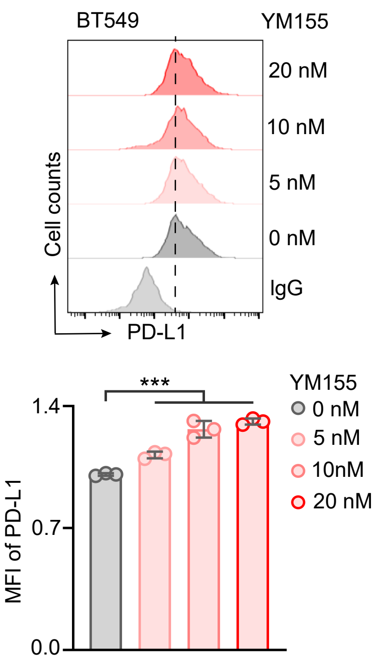


D

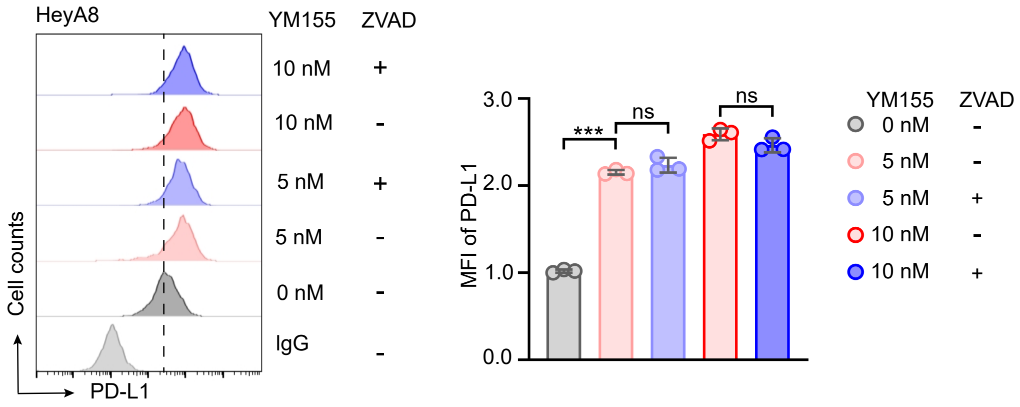

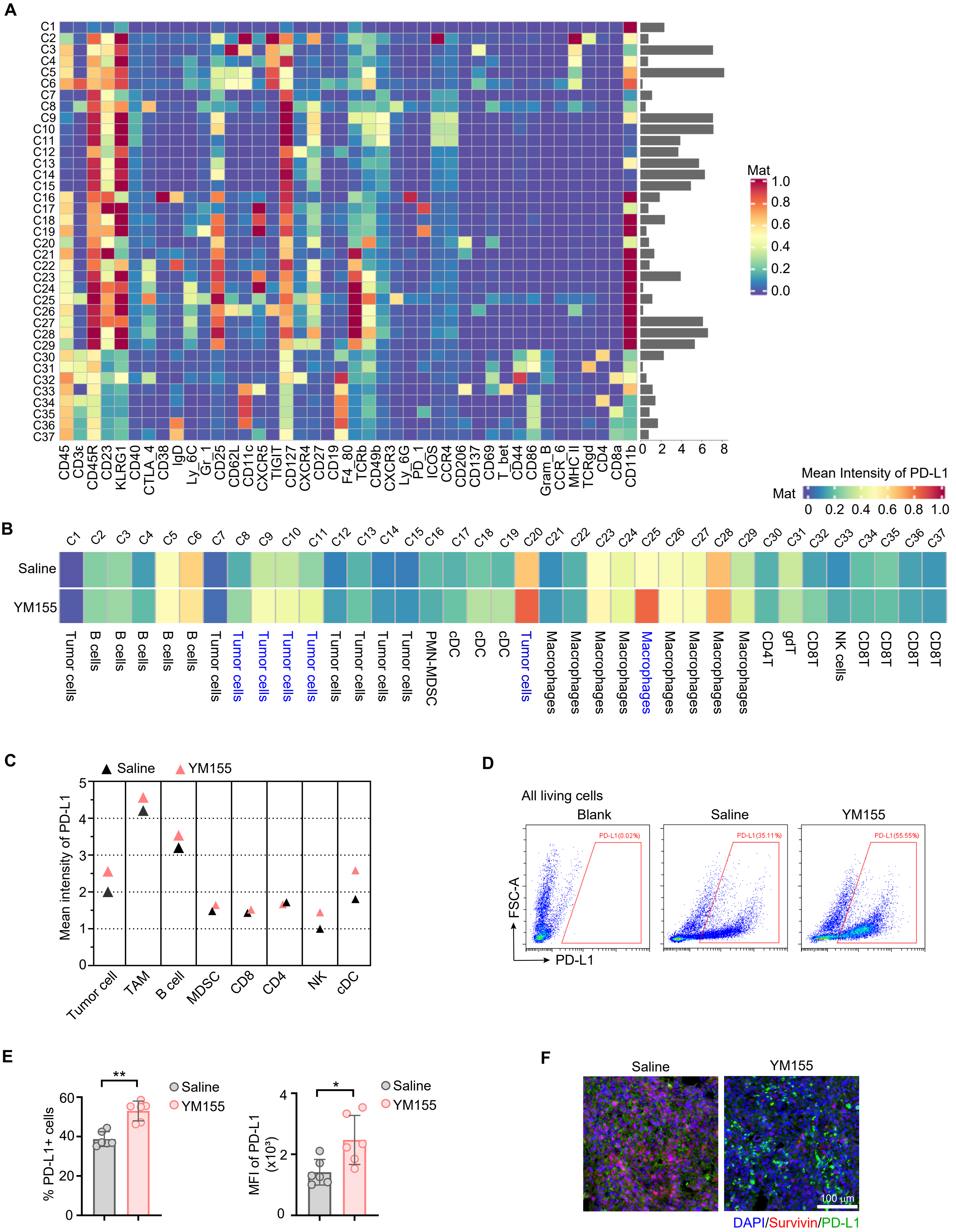

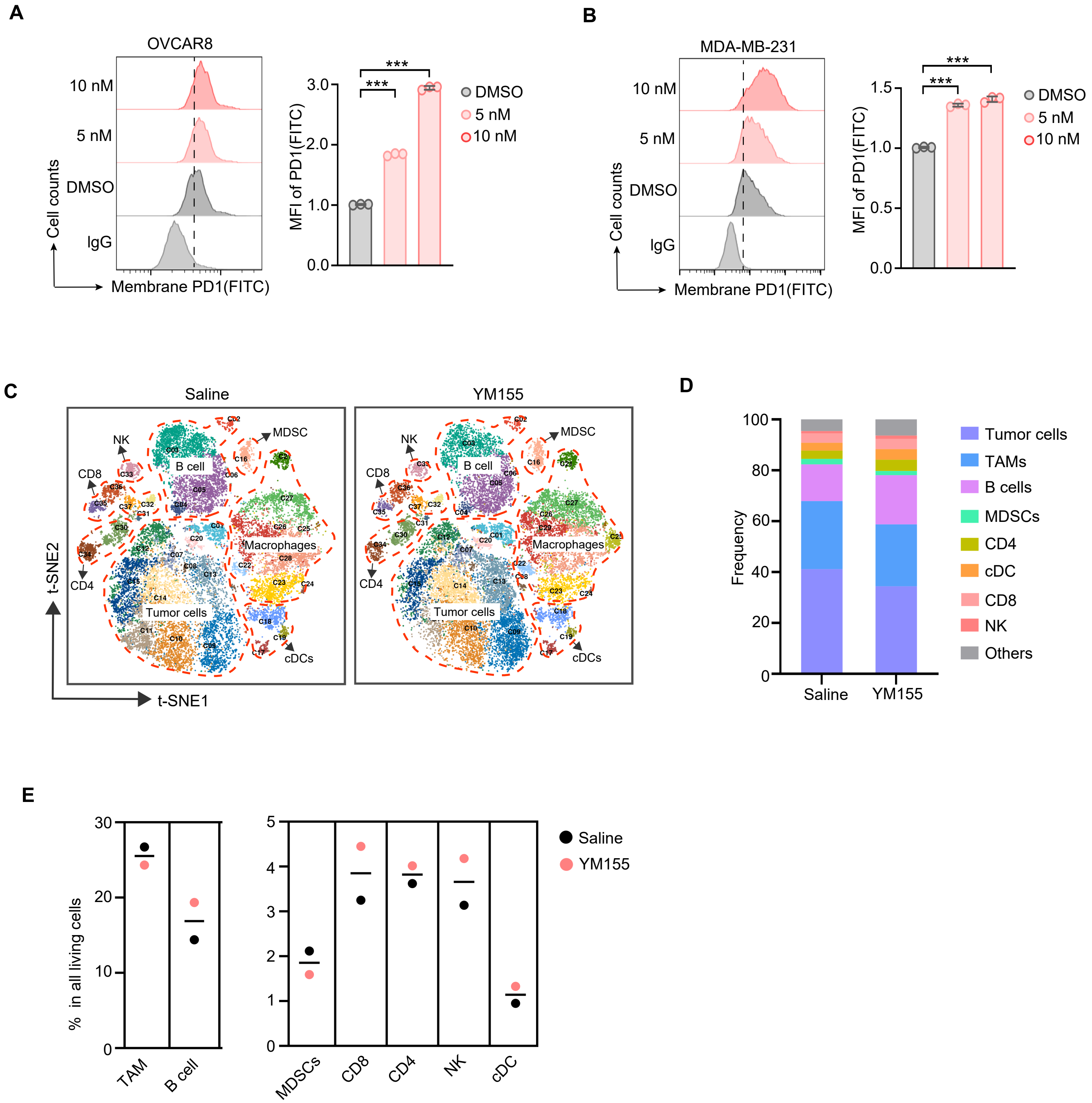

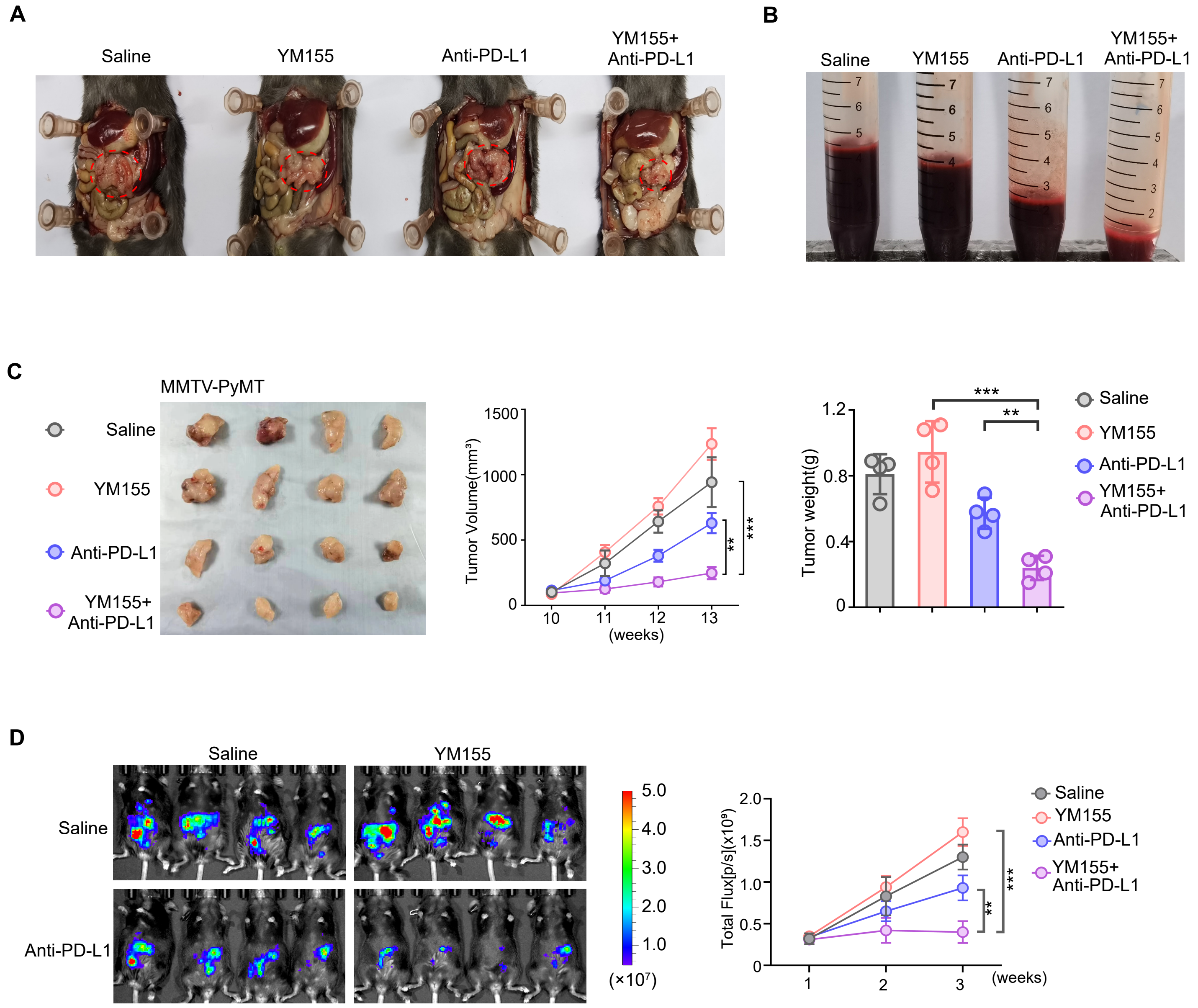

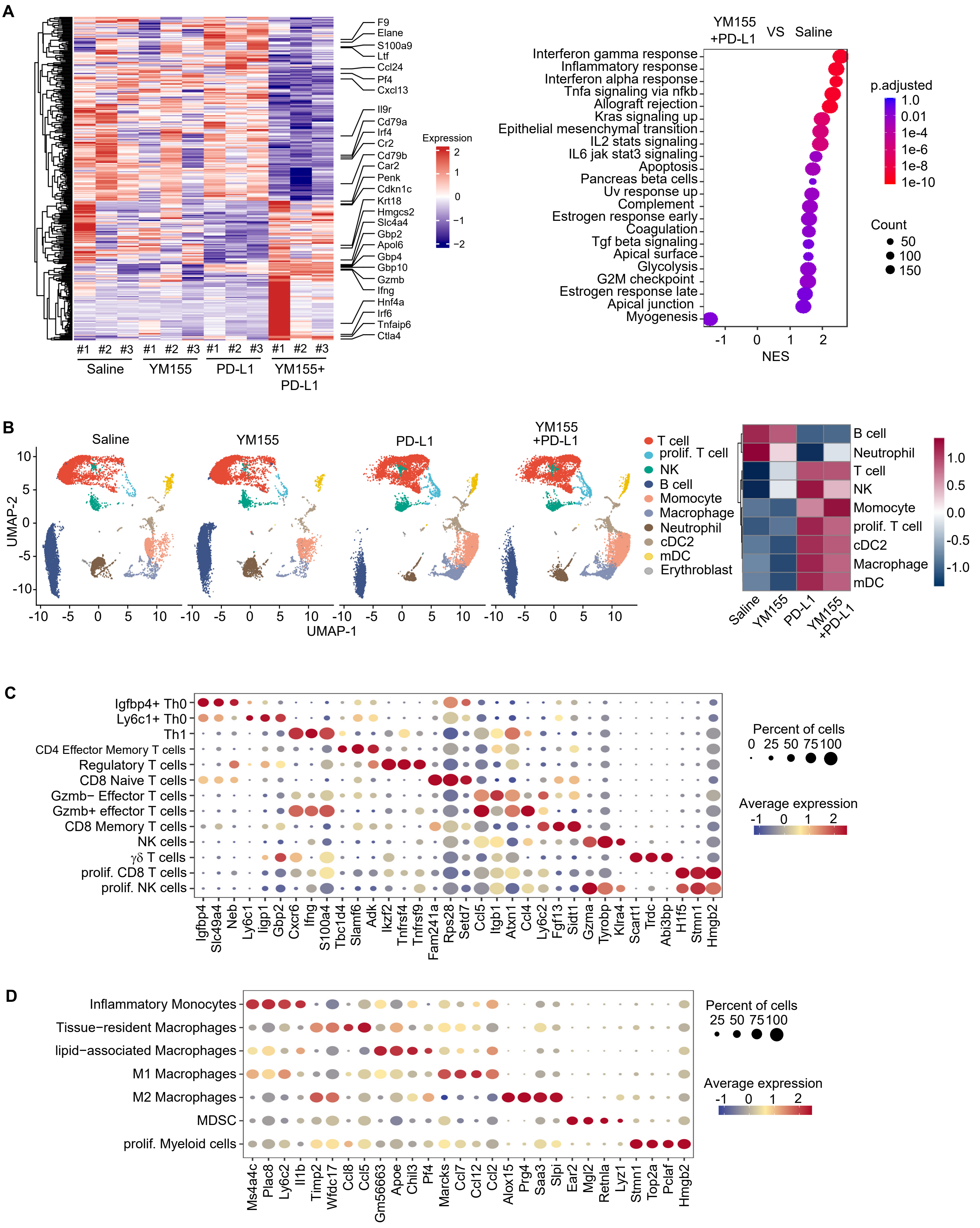

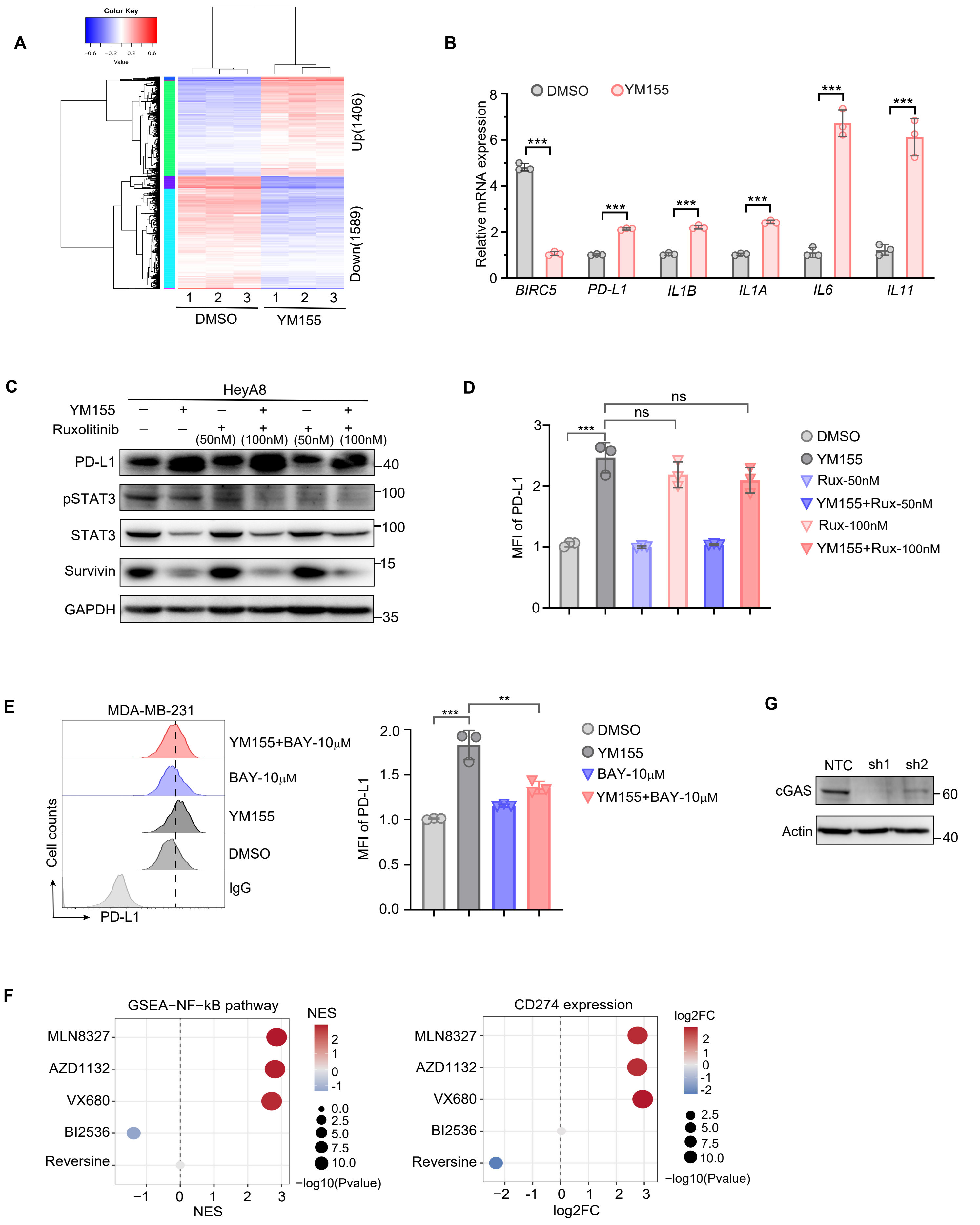

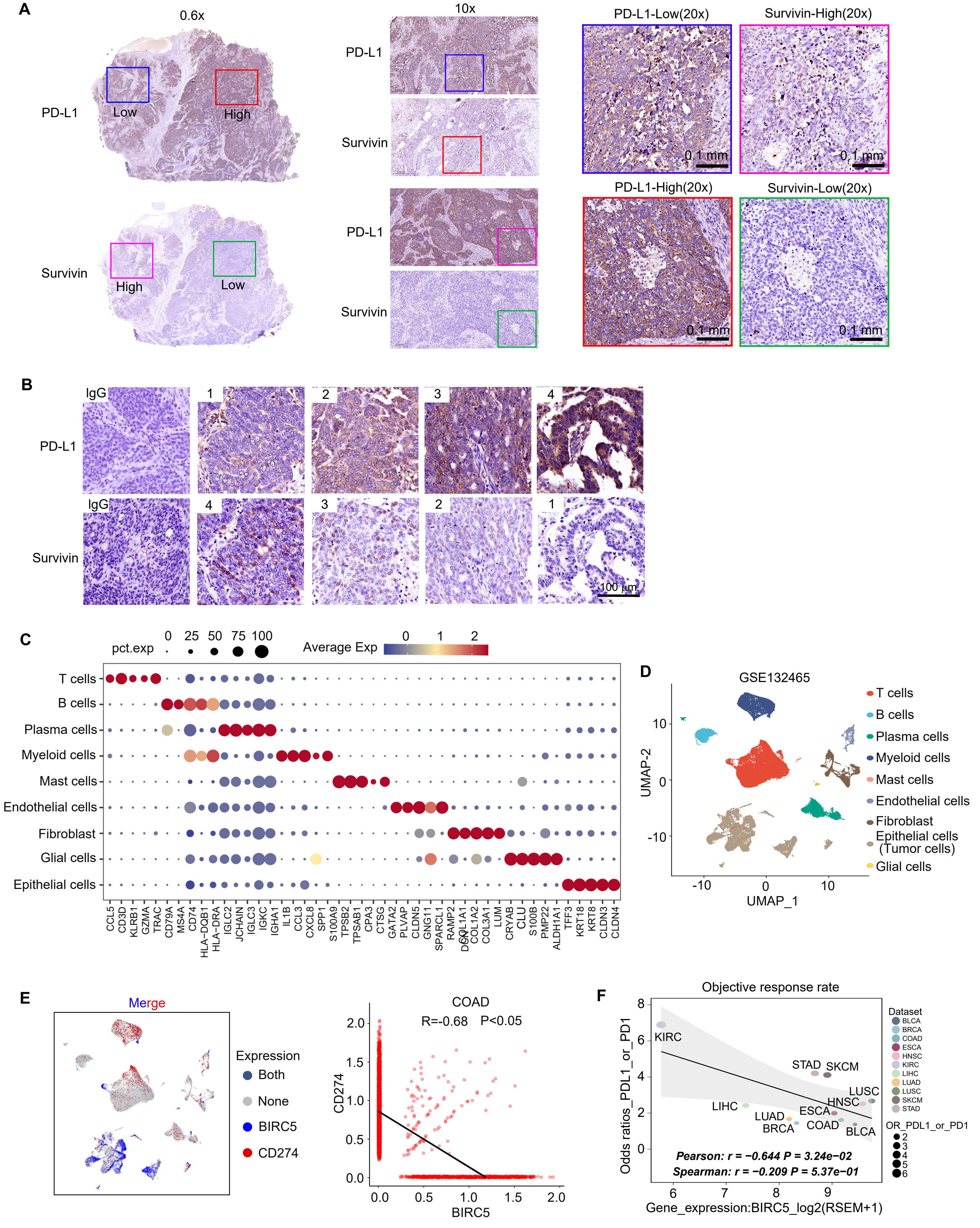
